# Transcriptomic, proteomic and functional consequences of codon usage bias in human cells during heterologous gene expression

**DOI:** 10.1101/2022.01.07.475042

**Authors:** Marion A.L. Picard, Fiona Leblay, Cécile Cassan, Anouk Willemsen, Josquin Daron, Frédérique Bauffe, Mathilde Decourcelle, Antonin Demange, Ignacio G. Bravo

**Affiliations:** Laboratory MIVEGEC (CNRS, IRD, University of Montpellier), French National Center for Scientific Research, Montpellier, France; BioCampus Montpellier (University of Montpellier, CNRS, INSERM), Montpellier, France

**Author notes:** Corresponding authors (MALP), (IGB). **AUTHOR CONTRIBUTIONS** Funding Acquisition, Project Administration and Supervision : IGB; Methodology : IGB, AD, MALP; Investigation : MALP, FL, CC, AW, FB, JD, MD, AD; Data Curation : MALP; Formal Analysis : MALP, IGB; Visualization : MALP; Conceptualization and Writing : MALP, IGB.

**Keywords:** Synonymous codon recoding, heterologous gene expression, translation, genotype to phenotype, mutation-selection

## Abstract

Differences in codon frequency between genomes, genes, or positions along a gene, modulate transcription and translation efficiency, leading to phenotypic and functional differences. Here, we present a multiscale analysis of the effects of synonymous codon recoding during heterologous gene expression in human cells, quantifying the phenotypic consequences of codon usage bias at different molecular and cellular levels, with an emphasis on translation elongation.

Six synonymous versions of an antibiotic resistance gene were generated, fused to a fluorescent reporter, and independently expressed in HEK293 cells. Multiscale phenotype was analysed by means of: quantitative transcriptome and proteome assessment, as proxies for gene expression; cellular fluorescence, as a proxy for single-cell level expression; and real-time cell proliferation in absence or presence of antibiotic, as a proxy for the cell fitness.

We show that differences in codon usage bias strongly impact the molecular and cellular phenotype: (i) they result in large differences in mRNA levels and in protein levels, leading to differences of over fifteen times in translation efficiency; (ii) they introduce unpredicted splicing events; (iii) they lead to reproducible phenotypic heterogeneity; and (iv) they lead to a trade-off between the benefit of antibiotic resistance and the burden of heterologous expression.

In human cells in culture, codon usage bias modulates gene expression by modifying mRNA availability and suitability for translation, leading to differences in protein levels and eventually eliciting functional phenotypic changes.

**IMPORTANCE:** Synonymous codons encode for the same amino acid, but they are not neutral regarding gene expression or protein synthesis. Bias between synonymous codons have evolved naturally and are also applied in biotechnology protein production. We have studied the multilevel impact of codon usage on a human cell system. We show that differences in codon usage lead to transcriptomic, proteomic and functional changes, modulating gene expression and cellular phenotype.

## INTRODUCTION

The canonical scenario of gene expression posits that DNA sequences are first transcribed into messenger RNA (mRNA) molecules that are secondly translated into proteins, so that one given nucleotide sequence encodes one predictable amino acid sequence ^1^. The initial version of this scenario did not provide any explanation on how a unique set of genes could be associated with several cellular phenotypes. Over the last decades, a large body of studies on gene expression have addressed this question and revealed multi-level regulation mechanisms increasing the diversity of the proteomic outputs that can be produced from a given genome. The standard genetic code that establishes a correspondence between the DNA coding units (*i*.*e*. the codon, a triplet of nucleotides, 64 in total) and the protein building blocks (*i*.*e*. the amino acids, 20 in total) is degenerated: 18 of the amino acids can individually be encoded by two, three, four or six codons, known as synonymous codons. In a first null hypothesis approach, one would expect synonymous codons to display similar frequencies. Instead, codon usage bias (CUBias, *i*.*e*. the uneven representation of synonymous codons ^2^) has been reported in a multiplicity of organisms, and varies not only between species but also within a given genome or even along positions in a gene ^3–8^.

The origin and the contribution of the different neutral and/or selective forces shaping CUBias constitute a classical research subject in evolutionary genetics. The hypothesis of translational selection proposes that differences in CUBias result in gene expression variations that ultimately lead to phenotypic differences, which could be subject to natural selection. Indeed, it has been established that variation in CUBias might constitute an additional layer of gene expression modulation ^9–11^. Notably, genetic engineering has extensively resorted to CUBias recoding for enhancing heterologous protein production, for its use in industrial applications or for vaccine design ^12–15^. The interaction between CUBias and the translation machinery has been well established, for instance in: (i) the co-variation of genomic CUBias and the tRNA content, from unicellular organisms ^4,16,17^ to metazoa (*Caenorhabditis elegans* ^18^, *Drosophila* ^19–21^, or humans ^22^; (ii) the correspondence between CUBias and expression level in bacteria ^23^ or in yeast ^24,25^; (iii) the increase in translation efficiency in bacteria when supplementing *in trans* with rare tRNAs ^26^; or (iv) the changes in tumorigenic phenotype in mice when switching from rare to common codons in the sequence of a cancer-related GTPase ^27^.

In contrast, a number of studies have communicated the lack of covariation between CUBias and gene expression (in bacteria, yeast, or human) ^28–31^; or even a negative impact of a presupposed “optimization”, which may in fact decrease the expression or the activity of the protein product ^32,33^. To address these conflicting results, it is important to tease apart the underlying mechanisms through which CUBias can impact the molecular, cellular and/or organismal phenotype. It has hitherto been established that CUBias can impact: (i) mRNA localisation, stability and decay ^34–38^ ; (ii) translation initiation ^31,39–41^ ; (iii) translation efficiency ^20,42–55^; and (iv) co-translational protein folding ^56–58^. But, fuelling the controversy, the respective contribution of each mechanism, if any, depends on the studied system, *e*.*g*. in which organism, whether the expressed gene is autologous or heterologous gene, or whether it has been recoded or not.

Finally, abundance and chemistry of transfer RNAs (tRNAs) introduces an additional layer of complexity (and thus an opportunity for regulation). In fast growing unicellular organisms, the tRNA gene content matches well codon usage preferences of the organism ^59^. Heterologous expression can thus be hampered by the lack or rarity of a tRNA that corresponds to a rare codon in the expression system of choice. Biotechnology engineering has circumvented this limitation by providing *in trans* the required tRNAs, encoding them into helper plasmids such as pRIG or pRARE ^60^. Further, many genomes actually do not contain dedicated tRNAs to decode each codon: *e*.*g*. for all amino acids encoded by two codons ending in C or U (Phe, Asn, Asp and His) the human genome contains only the tRNAs corresponding to the C-ending codons, which decode also the U-ending counterparts ^61^. Indeed, tRNAs are heavily modified and carry non-canonical nucleotides, which is often the case for an inosine residue in the first anticodon position ^62^, The non Watson-Crick base pairing interactions available to inosine allow to broaden codon-anticodon recognition ^63^, so that in bacteria and eukaryotes modified tRNAs carrying inosine in the anticodon can decode several synonymous codons for the amino acids Thr, Ala, Pro, Ser, Leu, Ile, Val and Arg ^64^. Thus, tRNA modification allows for one-to-many anticodon-to-codon translation potential, which may have had implications for the evolution of the protein repertoire in eukaryotes ^65^..

In this study, we aim at providing an integrated view of the molecular and cellular impact of alternative CUBias of a heterologous gene expressed in human cells. By combining transcriptomics, proteomics, fluorescence analysis and cell growth evaluation, we attempt to describe qualitatively, and to quantify as far as possible, the impact of CUBias and sequence composition of our focal heterologous gene on its own expression. These are usually called the *cis*-effects of CUBias on gene expression.

## RESULTS

### 1. Design of six synonymous gene versions that explore a large sequence space and cover a broad range of sequence composition variables

With the aim of analysing the effects of CUBias on protein synthesis, we have generated six synonymous variants of the gene encoding for the bleomycin-resistance protein from the bacterium *Streptoalloteichus hindustanus* (*shble*). We have chosen this heterologous protein as a reporter gene because it displays a mechanism of action (scission of DNA strands, ^66^) which is independent of the translation process, that we aim to study. For all six *shble* versions we added an AU1 tag in the N-terminus with the same nucleotide sequence, so that translation initiation will be similar for all *shble* versions and we can focus on the impact of CUBias on translation elongation. The *shble* ORFs were in-frame coupled via a P2A epitope to an *egfp* gene that encodes for a fluorescent protein reporter. The nucleotide sequence encoding for the P2A peptide was identical for all sequences and corresponds to that in the plasmid backbone. The expected heterologous transcript was a 1,602 base pair (bp) long mRNA encompassing a 1,182bp coding sequence (CDS). The CDS spanned the *AU1*-tag sequence in 5’, the *shble* bleomycin resistance reporter, the *P2A* peptide sequence inducing ribosomal skipping, and the *egfp* fluorescent reporter (Sup. Fig. 1). The presence of the AU1 epitope allowed us to use the same antibodies to detect the N-terminus of the SHBLE protein. The presence of the P2A peptide (NPGP motif) induces ribosome skipping ^67^, meaning that the ribosome does not perform the Gly-Pro transpeptidation bond and releases instead the AU1-SHBLE moiety and continues translation of the EGFP moiety.The synonymous versions of the *shble* were thus the only differences between constructs, and were characterized by distinct degrees of similarity to the average human CUBias (estimated using the COdon Usage Similarity INdex, COUSIN ^68^), GC composition at the third nucleotide of codons (GC3), and CpG dinucleotide frequency (CpG) (Table 1). Modifications in the *shble* sequence also entailed variations on the mRNA folding energy, calculated using the Vienna RNAfold webserver ^69^(Table 1). These four parameters combined allowed for a good discrimination of all constructs (Sup. Fig. 2), partly reflecting sequence similarities (Sup. Table 1). The COUSIN index quantifies the match between the CUBias of a focal sequence (in our case the different *shble* synonymous versions) and the chosen reference (in our case the average CUBias of human genes) ^68^. Briefly, the COUSIN is a normalised index so that a value of 1 corresponds to a focal sequence with similar CUBias to the reference; values above 1 correspond to similar CUBias to those in the reference, but of larger magnitude; a value of 0 corresponds to a lack of CUBias; and negative values correspond to CUBias opposite to those in the reference. The values for the COUSIN index exemplify the large variation in CUBias explored by our construct repertoire, ranging from “hyper-humanised” versions (namely shble#1 and shble#2, with COUSIN values above 2) to strongly “de-humanised” versions (namely shble#4 and shble#6, with negative COUSIN values).

**Table 1.** **Experimental conditions: the different constructs, and their sequence composition variables. Codon Usage Similarity index (COUSIN) values have been calculated against the average CUBias of human genes, as in** ^**68**^. **Folding energy values for the total mRNA transcripts have been calculated using the RNAfold Webserver** ^**69**^.

| Condition | Description | COUSIN of the *shble* sequence<sup>o</sup> | %GC3 of the *shble* sequence | %CpG of the *shble* sequence | Folding energy of the total transcript (kcal/mol) |
| --- | --- | --- | --- | --- | --- |
| **shble#1** | The most common codons in the human genome | 2.93 | 93.08 | 18.46 | -649.34 |
| **shble#2** | The GC-richest among the most common codons | 2.982 | 99.23 | 22.56 | -673.07 |
| **shble#3** | The AT-richest among the most common codons | -0.414 | 20.00 | 4.62 | -581.47 |
| **shble#4** | The rarest codons in the human genome | -1.651 | 33.85 | 20.51 | -613.49 |
| **shble#5** | The GC-richest among the rarest codons | 0.973 | 91.54 | 35.90 | -687.76 |
| **shble#6** | The AT-richest among the rarest codons | -0.924 | 9.23 | 0.51 | -543.50 |
| #empty | No *shble* but only *EGFP* CDS | n.a. | n.a. | n.a. | n.a. |
| #superempty | Neither *shble* nor *EGFP* CDS | n.a. | n.a. | n.a. | n.a. |
| mock | No plasmid | n.a. | n.a. | n.a. | n.a. |

### 2. Differences in CUBias of the *shble* gene resulted in differences in transcription

After transfection, we monitored DNA levels for transfection efficiency using quantitative PCR (qPCR) and we monitored mRNA levels for transcription efficiency using retrotranscription followed by qPCR (rt-qPCR) and RNA-sequencing (RNASeq). Analysis of the RNASeq read distribution revealed the presence of splicing events within the *shble* CDS for the two constructs with the lowest similarity to the human average CUBias, namely shble#4 (construct using the rarest codon for each amino acid) and shble#6 (using rare and AT-rich codons) (Sup. Fig. 3). The shble#6 transcript presented two spliced forms, using the same 5’ donor position and differing in three nucleotides at the 3’ acceptor position. The shble#4 transcript presented one spliced form, with donor and acceptor positions in the precise same locations observed for shble#6, despite the lack of identity in the intron-exon boundaries. The spliced intron (either 306 or 309 nucleotides long) was fully comprised within the 396 bp long *shble* sequence (Sup. Fig. 4), and the event did not involve any frameshift. Thus, *shble* splicing resulted in the ablation of the SHBLE protein coding potential without affecting the start codon and without modifying the EGFP coding potential. It is important to state that none of these alternative splicing events was predicted by the HSF (Human Splicing Finder) ^70^ nor the SPLM ^71^ splice detection algorithms used for sequence scanning during design. Analysis of mRNA abundances showed that the first spliced isoform (shared by both affected conditions) represented about 30% of the heterologous transcripts for shble#4, and 56% for shble#6. The second spliced isoform, exclusively found in condition shble#6, corresponded to 22% of the heterologous transcripts (Figure 1).

**Figure 1.**
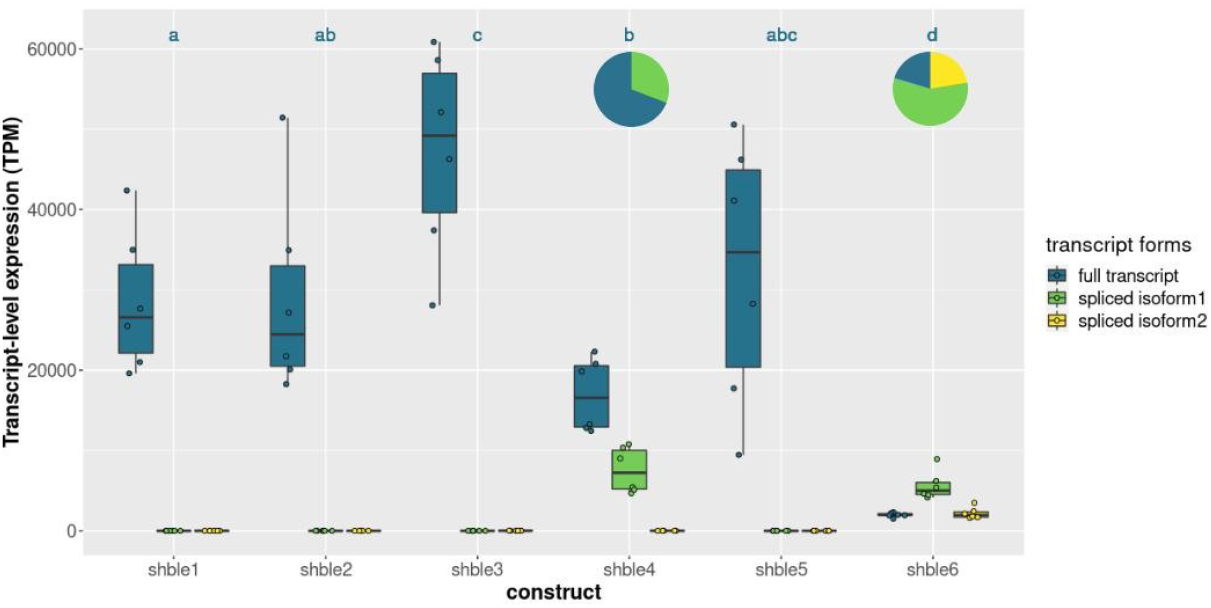
Transcript abundance after transfection with the different shble gene versions. mRNA-levels are expressed as transcripts per million values (TPM) for the full form (in dark blue) as well as for the two spliced forms (in green and yellow). Median values are given in Sup. Table 2. Pie charts illustrate the average proportions of the spliced forms detected in shble#4 and shble#6 conditions. The experiment was performed on six biological replicates. Dark blue letters above the different bars refer to the results of a Wilcoxon rank sum test. Conditions associated with a same letter do not display different median TPM values for the full mRNA (p<0.05 after Benjamini-Hochberg correction).

Full-length mRNA quantification showed differences in transcript levels across conditions, as follows: (i) the highest values were found in shble#3 (using the AT-richest among common codons); (ii) the variance was largest in shble#5 (using the GC-richest among rare codons); and (iii) shble#4 and shble#6 displayed the lowest mRNA abundance even when considering the sum of all isoforms (Figure 1, Sup. Table 2). We further verified that variations in transcript levels were not related to variations in transfection efficiency, by correcting the transcript levels after the plasmid DNA levels in each sample. After this normalisation, the above described pattern remained unchanged (Sup. Fig. 5). This suggests that variations in mRNA levels are not due to differences in the DNA level, and may instead be linked to the differentially recoded *shble* sequences.

In order to allow for further comparison between mRNA and protein levels, while accounting for the differential splice events, we have taken into account that the SHBLE protein was exclusively encoded by the full-length mRNA, while the EGFP protein could be translated from any of the three transcript isoforms. Hence, we used the ratio full-length mRNA over total heterologous transcripts (*i*.*e*. full-to-total ratio) to estimate the ratio of SHBLE-encoding over EGFP-encoding transcripts. This ratio was about 69% for shble#4, while for shble#6 it was close to 21% (Sup. Table 2). For the rest of the constructs, there was virtually no read corresponding to spliced transcripts and the ratio was in all cases above 99.96% (Sup. Table 2).

### 3. Differences in CUBias of the *shble* gene resulted in differences in SHBLE and EGFP protein levels

After transfection, and in paired samples with the mRNA analyses, we quantified SHBLE and EGFP protein levels by means of western-blot (Sup. Fig. 6, 7 and 8) and of label-free proteomics (Figure 2). Label-free proteomic analysis allowed to detect EGFP proteins for all constructs, with EGFP abundance in shble#3 and shble#6 being significantly lower than in other conditions (respectively 2.05 and 1.35 normalized iBAQ values, compared to an overall median of 10.08 for the other constructs) (Figure 2C, Sup. Table 3). The SHBLE protein was detected in all conditions but, for shble#6, it displayed extremely low abundance in five replicates and was not detected in one replicate (normalized iBAQ value of 0.03) (Figure 2B, Sup. Table 3). Further, the shble#3 condition displayed lower SHBLE protein levels than the remaining four other constructs (normalized iBAQ value of 0.93, compared to an overall median of 3.83) (Figure 2B, Sup. Table 3). Within a given condition, values for SHBLE and EGFP protein levels displayed a strong, positive correlation, albeit with a particular expression pattern for version shble#6 (Pearson’s R coefficients ranging from 0.82 to 0.95 depending on the condition; all p-values < 0.05; Figure 2A). The overall SHBLE-to-EGFP ratio was 0.46±0.1 for all constructs (ranging between 0.36 and 0.56 for the individual constructs), the exception being shble#6, which displayed very low ratio (0.03), as expected given the very low SHBLE levels (Figure 2D). For this specific construct, we find good correlation between SHBLE and EGFP levels (Pearson’s R = 0.93, p = 0.0072, Figure 2A), but the slope linking them is ten times lower than for any other construct (Figure 2D). Label-free proteomic quantification results were validated by image-based western blot quantification (Sup. Fig. 6, 7 and 8).

**Figure 2.**
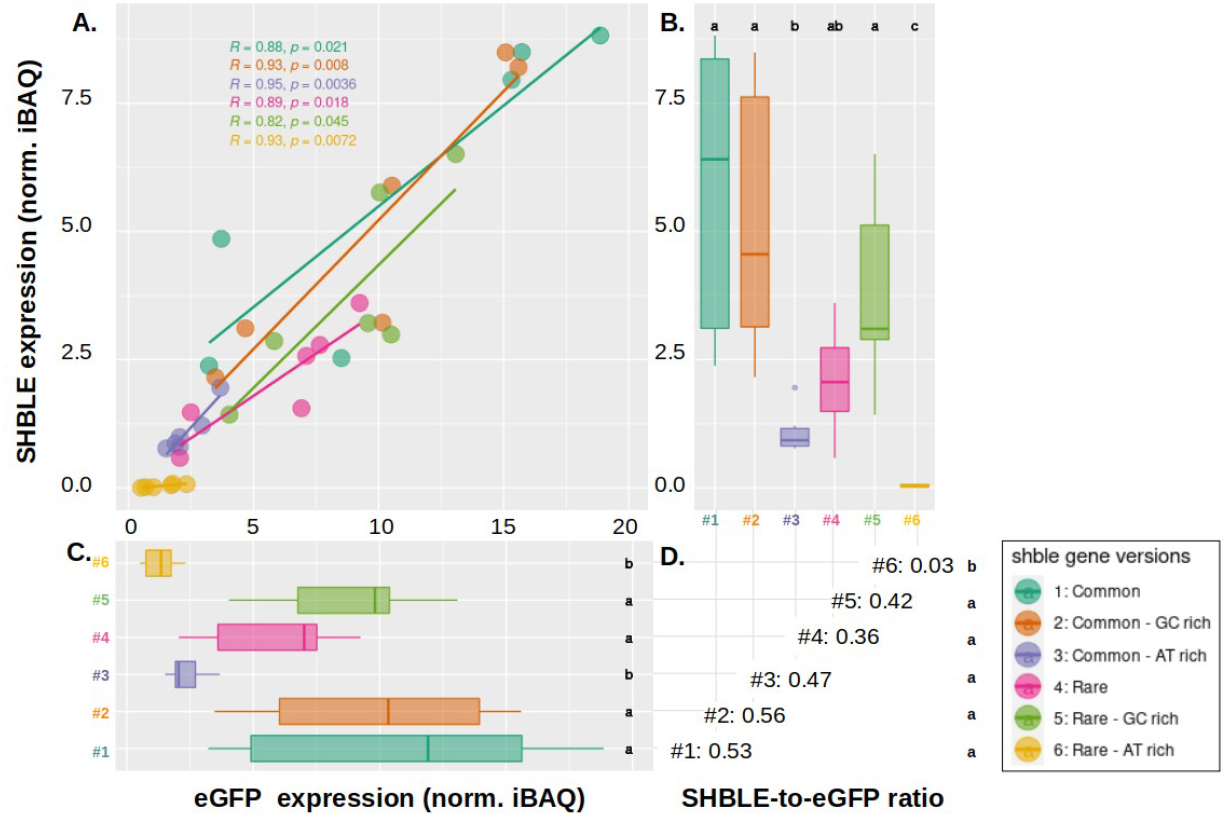
Expression of SHBLE and EGFP at the proteomic level, and relation between them. Panel A: Pearson’s correlation between SHBLE (y axis) and EGFP (x axis) protein levels. Six different conditions are shown: shble#1 (dark green), shble#2 (orange), shble#3 (purple), shble#4 (pink), shble#5 (light green) and shble#6 (yellow). Marginal boxplots (panels B and C) respectively show SHBLE and EGFP protein levels expressed as normalized iBAQ values. Median values are given in Sup. Table 3. The SHBLE-to-EGFP ratio for each of the six conditions (median of the ratios for each replicate) are given in panel D. Six replicates are shown (with three of them corresponding to two pooled biological replicates). Letters in the different panels refer to the results of a pairwise Wilcoxon rank sum test. Within each panel, conditions associated with a same letter do not display different median values of the corresponding variable (p<0.05 after Benjamini-Hochberg correction).

### 4. Differences in CUBias and mRNA physicochemical properties partly explain differences in translation efficiency

After separately analysing mRNA and protein levels in cells transfected with the different *shble* versions, we aimed at establishing a connection between transcription and translation levels. Because SHBLE and EGFP protein analyses led to similar results, we focus here only on SHBLE. We chose to normalise the protein levels over the corresponding mRNA levels, and we interpret this protein-to-mRNA ratio as a proxy for translation efficiency. The median values of the protein-to-mRNA ratio were similar for constructs #1, #2, #4 and #5, whereas conditions #3 and #6 were discordant (Figure 3A and Sup. Table 4): translation efficiency is over five times lower for condition #3 (which displayed high transcription levels) and over thirteen times lower for condition #6 (Sup. Table 4). Overall, variation in full-length transcript levels explained 45% of the variation in SHBLE protein levels (Pearson’s R=0.45, p = 0.0054) (Sup. Fig. 9). This explanatory power of mRNA levels over protein levels increased to 68% when considering only conditions #1, #2, #4 and #5 (Pearson’s R=0.68, p = 0.00025). As discussed below, these values fit well in previous descriptions in the literature about the explanatory power of variations at the mRNA level to account for variations at the protein level for eukaryotic cells ^72^.

**Figure 3.**
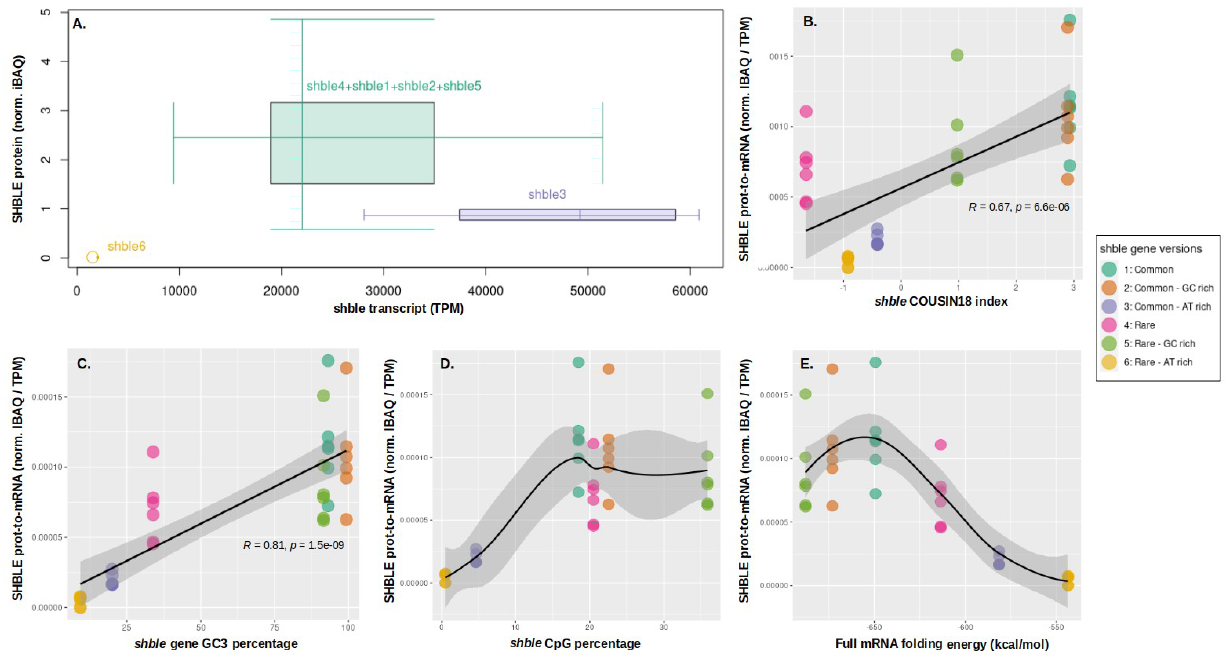
Variation of translation efficiency as a function of CUBias and mRNA physicochemical parameters. **Panel A:** Combined distribution of SHBLE protein level (y axis - normalized iBAQ) and shble transcript level (x axis - TPM); individual construct boxes are condensed in a single one when the squares defined by the first and third quartiles overlaps (which is the case for shble#1, shble#2, shble#4 and shble#5, shown condensed in dark green). For each construct, median values are given in Sup. Table 4. **Panel B:** Pearson’s correlation between SHBLE protein-to-mRNA ratio and COUSIN index of the shble recoded version. **Panel C:** Pearson’s correlation between the SHBLE protein-to-mRNA ratio and the GC3 percentage of the shble recoded version. **Panel D:** SHBLE protein-to-mRNA ratio variations depending on CpG frequency of the shble recoded version. **Panel E:** Correspondence between the SHBLE protein-to-mRNA ratio and the folding energy of the corresponding transcript. Curves in panels D and E correspond to a LOWESS (LOcally WEighted Scatter-plot Smoother) local polynomial regression to visually display co-variation between the two variables plotted. The results for six full biological replicates are shown, each of them with independent RNAseq measurements but pooled by pairs for the label-free proteomic analysis.

In order to understand the differential translation efficiency between constructs, we explored the explanatory potential of four sequence composition and mRNA physicochemical parameters. We observe first that the closer the CUBias of the *shble* synonymous versions to the average human CUBias, the higher the translation efficiency in our human cells in culture (Pearson’s R=0.67, p=6.6e-6, Figure 3B). The exception to this trend was condition shble#4 which displayed the lowest match to the human CUBias, but a higher protein-to-transcript ratio than shble#6 or shble#3 (Figure 3B). The lower ratio for these two later conditions could be explained at the light of the three other tested parameters. Indeed, an increase of GC3 content corresponded monotonically to an increase in the protein-to-transcript ratio (Pearson’s R=0.81, p=1.5e-9, Figure 3C), and shble#6 and shble#3 had the lowest GC3 content. Increase in CpG frequency (Figure 3D) resulted in an increased translation efficiency that reached a plateau for all recoded forms beyond 20% CpG presence, even for the very CpG-rich form shble#5. Finally, variation in mRNA folding energy (Figure 3E), corresponded to a bell-shaped variation in SHBLE protein-to-transcript ratio so that both low and high values resulted in decreased translation efficiency. Thus, the shble#3 condition combined suboptimal values for all four studied characteristics and resulted in poorly efficient translation in spite of the high mRNA levels (see part 2). In contrast, shble#1 (recoded using the most used codons), displayed maximum values for each parameter, thus resulting in the most efficient translation (highest protein-to-mRNA ratio).

### 5. Differences in CUBias lead to differences in EGFP protein expression at population level, but also at single-cell level

We have demonstrated above that variation in SHBLE protein levels were highly correlated to variation in the EGFP fluorescent reporter (Figure 2A). On this basis, and in order to further assess the phenotypic variation at the single-cell level, we performed an extensive analysis of the cell-based fluorescence values of 16 transfection replicates by means of fluorescent cytometry analyses. We verified first that the total fluorescence signal (*i*.*e*. the total fluorescence levels in the cell population) was strongly correlated to the EGFP level estimated by the label-free proteomics (Pearson’s R=0.86, p=4.8e-15, Sup. Fig. 10). We observed then that the single-cell distribution of this fluorescence signal was (i) different for all conditions from that obtained with cells expressing EGFP alone (*i*.*e*. “empty” control; individual Anderson-Darling test results shown in Table 2); and (ii) multimodal for all the conditions expressing EGFP (Figure 4A, Sup. Fig. 11). We have approximated these multimodal cell populations by means of curve deconvolution, and showed that a composite distribution based on two underlying Gaussian-like cell populations fitted well the observed distributions (Sup. Fig. 12). We conclude thus that synonymous variation of the upstream *shble* sequence modulated and modified the individual cell fluorescence phenotype, and that for a given version of the *shble* sequence, cells were differentially impacted by the construct expression, overall defining two subpopulations of low or high EGFP expression.

**Table 2.**
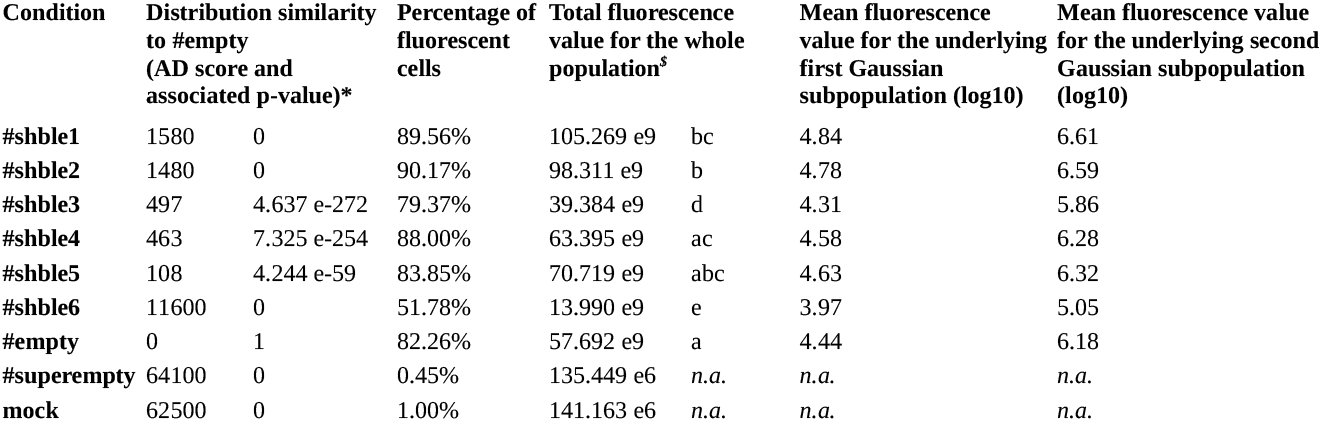
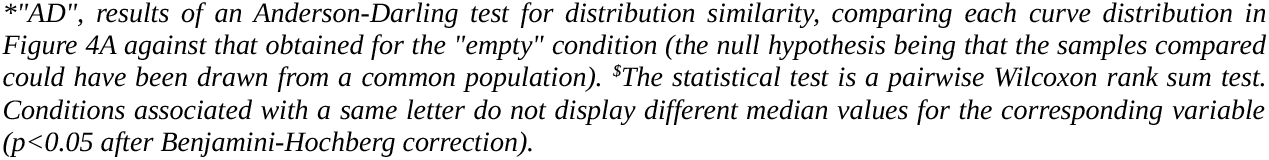
Quantitative parameters of green fluorescence signal distribution per condition.

| Condition | Distribution similarity to #empty (AD score and associated p-value)* |  | Percentage of fluorescent cells | Total fluorescence value for the whole population<sup>§</sup> |  | Mean fluorescence value for the underlying first Gaussian subpopulation (log10) | Mean fluorescence value for the underlying second Gaussian subpopulation (log10) |
| --- | --- | --- | --- | --- | --- | --- | --- |
| #shble1 | 1580 | 0 | 89.56% | 105.269 e9 | bc | 4.84 | 6.61 |
| #shble2 | 1480 | 0 | 90.17% | 98.311 e9 | b | 4.78 | 6.59 |
| #shble3 | 497 | 4.637 e-272 | 79.37% | 39.384 e9 | d | 4.31 | 5.86 |
| #shble4 | 463 | 7.325 e-254 | 88.00% | 63.395 e9 | ac | 4.58 | 6.28 |
| #shble5 | 108 | 4.244 e-59 | 83.85% | 70.719 e9 | abc | 4.63 | 6.32 |
| #shble6 | 11600 | 0 | 51.78% | 13.990 e9 | e | 3.97 | 5.05 |
| #empty | 0 | 1 | 82.26% | 57.692 e9 | a | 4.44 | 6.18 |
| #superempty | 64100 | 0 | 0.45% | 135.449 e6 | n.a. | n.a. | n.a. |
| mock | 62500 | 0 | 1.00% | 141.163 e6 | n.a. | n.a. | n.a. |

*"AD", results of an Anderson-Darling test for distribution similarity, comparing each curve distribution in* *Figure 4A against that obtained for the "empty" condition (the null hypothesis being that the samples compared* *could have been drawn from a common population). <sup>\$</sup>The statistical test is a pairwise Wilcoxon rank sum test.* *Conditions associated with a same letter do not display different median values for the corresponding variable* *( $p < 0.05$ after Benjamini-Hochberg correction).*

**Figure 4.**
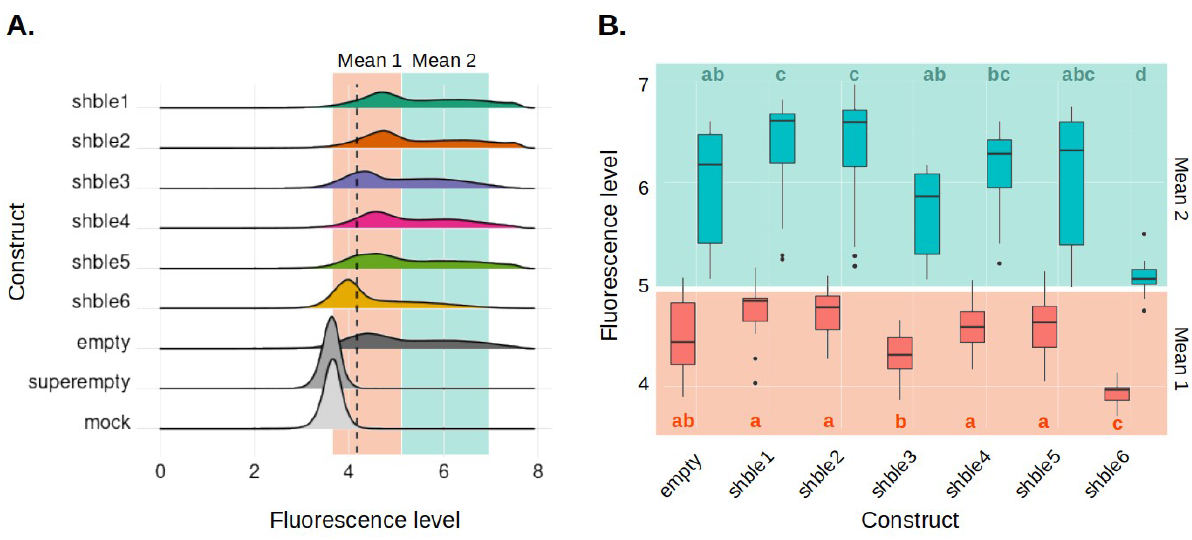
Distribution of the fluorescence signal for the different constructs, and mean values of the two gaussian curves modeling the fluorescence distribution. Panel A depicts the density of the green fluorescence signal (log10(FITC-A)) considering 480,000 individual cells for each condition: shble#1 (most common codons, dark green), shble#2 (common and GC-rich codons, orange), shble#3 (common and AT-rich codons, purple), shble#4 (rarest codons, pink), shble#5 (rare and GC-rich codons, orange light green), shble#6 (rare and AT-rich codons, yellow). The positive control is “empty” (i.e. transfected cells, expressing EGFP without expressing SHBLE, in dark grey); and the negative controls are “superempty” (i.e. transfected cells, not expressing EGFP nor SHBLE, in medium grey) and “mock” (i.e. untransfected cells, in light grey). The dashed black line shows the threshold for positivity (14,453 green fluorescence units, corresponding to 4.16 in a log10 scale). Panel B represents the first gaussian mean1 (population of lower intensity, in red), and the mean2 (population of higher fluorescence intensity, in blue). Values in the y-axis (cell fluorescence) are continuous, and the red and blue colours are for representation purposes only. For each category (mean1 and mean 2), the statistical test is a pairwise Wilcoxon rank sum test, with Benjamini-Hochberg adjusted p-values on sixteen biological replicates: for each colour, conditions associated to a same letter do not display different median values of the corresponding variable.

For each condition, we describe the fluorescence behaviour of the whole cell population using the following summary statistics (Table2): (i) the fraction of cells displaying fluorescence above the cell autofluorescence threshold (discontinuous line in Figure 4A); (ii) the total fluorescence value of the whole population; (iii) the median fluorescence value of the population; (iv) the mean fluorescence value for each underlying Gaussian populations. We observed that the median fluorescence value of the population correlated very well with the overall fluorescence (R=0.85, p-value<2.2e-16, Sup. Fig. 13), but that the later allowed for a better discrimination between conditions. Conditions shble#1 and shble#2 displayed the highest fluorescence values, while shble#3 displayed ca. 2.5 times lower fluorescence values and shble#6 over seven times lower fluorescence values (Table 2, Sup. Fig. 13). Differences in total fluorescence levels reflected a reproducible impact of the synonymous construct expression on the complete cell population, independently of whether individual cells displayed very high or very low fluorescence: indeed, between each condition, both underlying Gaussian curves shifted following the same pattern, as illustrated by the variations of their mean values (Figure 4B, Table 2). When combining all our summary statistic variables into a principal component analysis for describing the cellular population fluorescence we observed that indeed shble#6, and to a lesser extent shble#3, were the most divergent conditions, characterized by the highest proportion of negative or low-fluorescent cells, while shble#1 and shble#2 displayed very similar behaviour characterized by high fluorescence values in all scores (Sup. Fig. 14). These results strengthened the observations obtained by the label-free proteomic experiments, and underlied the cell-to-cell reproducibility of the impact of synonymous substitutions.

### 5. Differences in CUBias of the *shble* gene resulted in differences in cell growth dynamics and antibiotic resistance

Finally, since our *shble* reporter gene confers resistance to the bleomycin antibiotic, we aimed at quantifying the functional impact of the different molecular phenotypes described above on cellular fitness. For this, we performed a real-time cell growth analysis, both in presence and in absence of antibiotics. For all conditions, we monitored over time a dimensionless parameter named “Cell Index”, that integrates cell density, adhesion, morphology and viability; and we evaluated the total area below the curve as a proxy for cell growth (Sup. Method 2.8). We fitted to a Hill’s equation the variation of Cell Index values (*i*.*e*. cell growth) as a function of the antibiotic concentration, so that we could recover for each condition: (i) the maximum growth in the absence of antibiotic (Figure 5, variable for the y axis); and (ii) the estimation for the antibiotic concentration value that inhibited cell growth to half the maximum (IC50; Figure 5, variable for the x axis). Higher values of the variable “maximum growth in the absence of antibiotics” reflect a lower impact of the heterologous construct in the cell, while higher values of the IC50 variable reflect a higher resistance potential of the cell to face the bleomycin antibiotic. The results show that cell populations that grow more in the absence of antibiotics correspond also to cell populations that resist higher antibiotic concentrations. We interpret that this connection between growth variables reflects a trade-off between (i) the potential benefit of the antibiotic resistance - conferred by SHBLE expression, and realised only in the presence of antibiotics-, and (ii) the cost incurred through heterologous protein overexpression -associated to both SHBLE and EGFP expression and that is present independently of the presence of the antibiotic. This trade-off results indeed in a non-monotonic relationship between heterologous protein levels and cell growth: condition shble#6 produces low levels of heterologous protein and thus allows for the highest growth in the absence of antibiotics, but it does not confer the highest resistance levels; while conditions shble#1 and shble#2 produce the highest amounts of heterologous proteins and incur thus in a substantial burden, heavier than the potential benefit of the conferred antibiotic resistance (Figure 5A, and Sup. Fig 15).

**Figure 5.**
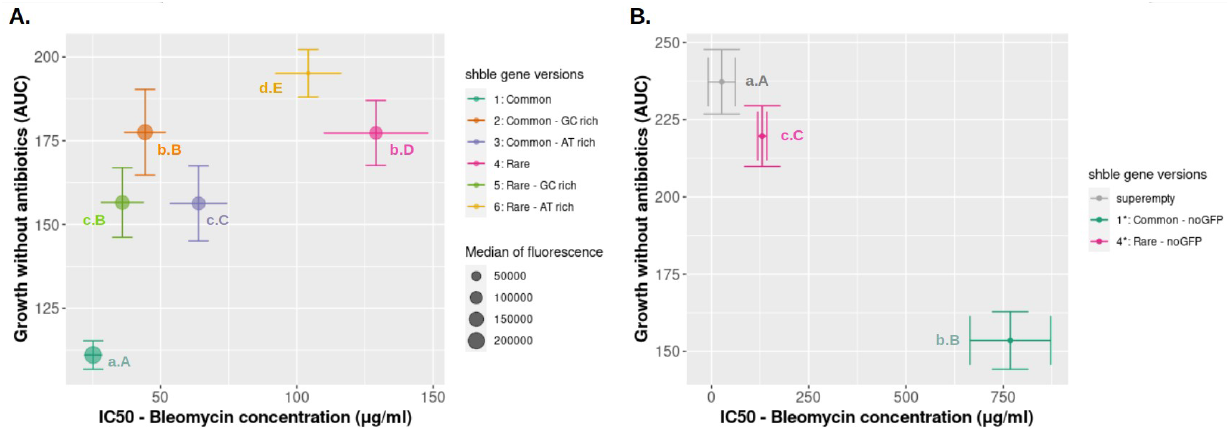
Variation of cell growth in presence or in absence of antibiotics, for gfp-coupled constructs (A) or gfp-free constructs (B). For both panels, the y axis represents maximum cell growth in absence of antibiotics, proxied as the area under the curve of the delta Cell Index (“AUC”); and the x axis represents the bleomycin concentration that reduces to 50% the corresponding growth (“IC50”). The plotted central values were estimated fitting variation of Cell Index data to Hill’s equation (pooled data, 3 to 6 biological replicates), and bars correspond to the estimated standard error. Statistical tests are Welch modified two-sample t-tests, performed for the AUC (small letters, y axis) or the IC50 (big letters, x axis): for each size of letters, conditions associated with a same letter do not display different median values of the corresponding variable (p<0.05 after Benjamini-Hochberg correction). As an example for interpretation, orange and pink values are not different in the y-axis projection (labelled both with b) but differ on their x-axis projection (labelled respectively with B and D). For panel A, the size of the dots is proportional to the corresponding total of fluorescence, which is used as a proxy for the level of heterologous proteins. Six different conditions are shown: shble#1 (dark green), shble#2 (orange), shble#3 (purple), shble#4 (pink), shble#5 (light green), shble#6 (yellow). For panel B, three different conditions are shown: superempty control (grey), versions shble#1* (dark green) and shble#4* (pink) lacking the EGFP reporter gene.

We aimed at disentangling the two forces in this trade-off by testing two additional constructs containing solely versions shble#1 and shble#4 of the *shble* gene, and not linked to the *egfp* reporter (labelled #1* and #4* in Figure 5B). Comparing the growth-related variables for *shble* versions #1 and #4 with or without EGFP, both versions shble#1* and shble#4* displayed a similar increase in max. growth in the absence of antibiotic with respect to their EGFP+ relative counterparts (respectively 38% and 24%). However, while the IC50 of shble#4* remained similar to shble#4, the antibiotic resistance for version shble#1* dramatically increased with respect to that of shble#1. Notwithstanding, in the absence of antibiotic shble#4* still performed better than shble#1*.

Overall, we interpret that (i) in the absence of antibiotic, higher amount of heterologous protein (independently of whether they correspond to SHBLE, or to SHBLE and EGFP) had pronounced negative impact on cell fitness; and that (ii) in the presence of antibiotic, the optimum between the conferred resistance and the cost of protein burden was determined by both, the total amount of heterologous proteins, and the abundance of the SHBLE protein, conferring antibiotic resistance.

## DISCUSSION

In the present manuscript we have analysed the multilevel molecular effects of CUBias on gene expression and have further explored higher-level integration consequences at the cellular level. We have focused on the effects of CUBias of our focal *shble* gene on its own expression and function, *i*.*e*. the so-called *cis*-effects of CUBias. The global *trans*-effects of CUBias of our focal gene on the expression levels of other cellular genes have been analysed and described in an accompanying manuscript ^73^. Our results show that a combination of synonymous changes results in important multilevel variation in gene expression levels and leads to dramatic differences in the cellular phenotype. We summarize our observations of these *cis*-effects in Figure 6, which displays variation in each of the variables that we have monitored, either experimentally or sequence-dependent. Conditions shble#1 and shble#2 display a very similar global profile, consistent with the fact that they are identical in 95.6% of their sequence (Table S1). Further, conditions with CUBias close to human average one (shble#1, shble#2 and shble#5) cover a similar phenotypic space, very different from the phenotypic space covered by conditions with CUBias opposite to the human average (shble#3, shble#4 and shble#6).

**Figure 6.**
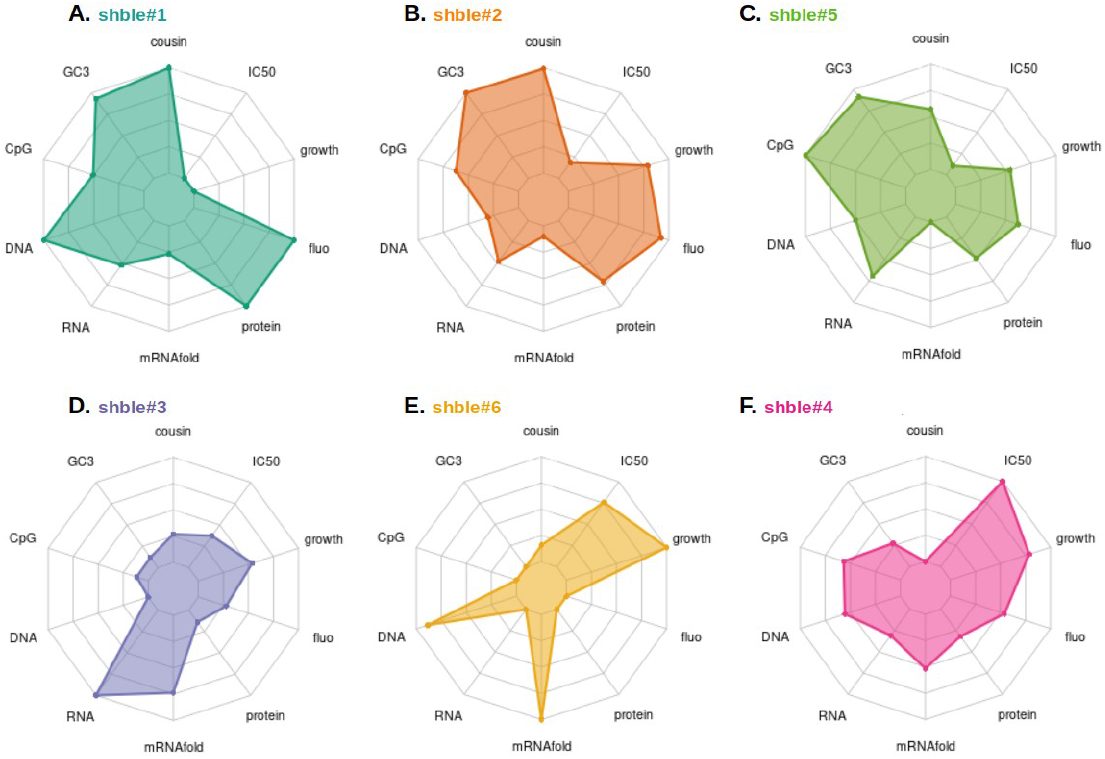
Summarizing combination of sequence composition parameters and multi-level phenotypes for each customized version of the shble antibiotic resistance gene. The six versions, designed with the one amino acid – one codon strategy, are shown by decreasing similarity to the average genome human CUBias (i.e. expressed as their cousin score, (59)): CUBias for shble#1 and shble#2 are similar to the human CUBias but of larger magnitude, #5 CUBias is similar to the human CUBias, and #3, #6, #4 CUBiases are opposite to the human CUBias. **A**. shble#1 (most common codons, in dark green), **B**. shble#2 (common and GC-rich codons, in orange), **C**. shble#5 (rare and GC-rich codons, in light green), **D**. shble#3 (common and AT-rich codons, in purple), **E**. shble#6 (rare and AT-rich codons, yellow) and **F**. shble#4 (rarest codons, in pink). The sequence characteristics are from the top to the left: “cousin”; “CpG” (the CG dinucleotide frequency in the recoded shble), “GC3” (the GC content at the third codon position in the recoded shbl), and “mRNAfold” (the mRNA folding energy for the recoded shble transcript). The different phenotypic variables, from the bottom to the right are: “DNA” (the transfection efficiency, estimated represented by the amount of plasmid after qPCR), “RNA” (the amount of SHBLE-coding full mRNA, estimated after rt-qPCR), “protein” (the amount of SHBLE protein, estimated by quantitative proteomics), “fluo” (the total fluorescence signal, estimated by flow cytometry), “growth” (proxy of the cellular fitness in absence of antibiotics, estimated by real-time cell growth analysis) and “IC50” (proxy of the cellular fitness in presence of antibiotics, estimated by real-time cell growth analysis). All variables have been independently re-scaled for representation purposes, from lowest (central) to highest (periphery) value.

### Variation in CUBias modifies alternative splice patterns

In the heterologous transcripts for the two versions with the most dissimilar CUBias with respect to the human average (shble#4 and shble#6) we identified splicing events, located within the *shble* ORF, that were not detected by leading splice site predicting algorithms ^70,71^. Splicing ablated the SHBLE coding potential without modification of the EGFP coding potential. The spliced transcripts amounted to 20% and 80% of all heterologous transcripts in shble#4 and shble#6 respectively. Variation in CUBias across intron-exon boundaries has been described in several eukaryotes (*e*.*g*. human, fishes, fruit flies, nematodes, plants ^11,74,75^); and splicing regulatory motifs that can be disrupted by synonymous mutations have been described in mammals ^9,75–78^. A reduced single nucleotide polymorphism density and a decreased rate of synonymous substitutions have further been reported in these regulatory regions, which can be interpreted as a signature for selective pressure ^79,80^. Thus, selection against mRNA mis-processing can constitute an important selective force that results in concomitant selection for a precise local CUBias ^81^. This selective force has even been proposed to outperform translational selection in *Drosophila melanogaster* ^82^. It is interesting to state here that we did not detect any western blot signal in any of our nine biological replicates that could correspond to the spliced, short SHBLE polypeptides in the shble#4 and shble#6 conditions (see Sup. Figs. 6, 7 and 8). Further, we did not detect in our proteomic analyses any trace of the expected peptides that could differentiate the spliced SHBLE versions from the full length SHBLE protein. For western-blot detection we used an AU1 epitope located in the N-terminus of the protein, and that should be present in all SHBLE forms, spliced or not. Lack of western-blot detection of these N-terminal spliced short SHBLE polypeptides could simply reflect a technical limitation, as they are barely 54 amino acids-long (or 53 for the minor spliced version in shble#6). However, in the case of shble#6 the lack of concordance between SHBLE levels and EGFP levels (see the very low slope in Figure 2A and 2B) suggests rather a genuine very low presence of spliced SHBLE molecules in our samples. We interpret that our results are rather compatible with a low stability of the spliced, short SHBLE versions, possibly linked to a faulty folding that could lead to a rapid degradation upon synthesis. Indeed, these spliced, short SHBLE versions span less than 20 amino acids of the original SHBLE sequence, so that the protein that is expected to fold into the known quaternary structure ^83^ actually does not exist after splicing.

### Variation in CUBias correlates with differences in mRNA levels

We observe significant differences in mRNA levels among the recoded *shble* versions, with shble#3 showing over five times more full-length mRNA than shble#6 (Table S2). Transcript abundance at a given time point is the result of integrating mRNA synthesis and degradation kinetics. In our experimental setup differential transfection efficiency does not explain differences in mRNA levels because variation in mRNA levels between conditions was independent of variation in DNA abundance. We interpret as well that differential ribosomal recruitment is unlikely to explain differences in mRNA levels, all our synonymous constructs share the same CMV promoter, the 5’ untranslated region, and the AUG context. We interpret therefore that the observed differences in mRNA levels probably result from differential mRNA stability and decay, rather than from primary transcription regulation. Such an effect has been described for bacteria (*E. coli* ^84^), unicellular eukaryotes (*S. cerevisae, S. pombe* ^35^, *N. crassa, T. brucei* ^*85,86*^), and metazoa (fruit fly ^87^ or zebrafish ^88^). In human cells, it has been shown that nucleotide composition and CUBias have an impact on mRNA stability, so that transcripts with longer half-lives are enriched in GC-rich codons ^89^, possibly through translation-associated decay mechanisms ^90^. Nevertheless, in our experimental design heterologous mRNA levels are not a monotonic function of mRNA composition, as versions shble#3 and shble#6 are the AT-richer ones (respectively 20% and 10% GC3) but display respectively the highest and the lowest levels of heterologous mRNAs. The effects on version shble#6 are difficult to address as only 20% of the total heterologous transcripts contain the customized *shble* sequence. It is thus impossible to disentangle the effects of sequence composition on the full mRNA level from the consequences of the splicing defect. The very high transcript levels and very low protein levels for version shble#3 are interesting in the light of recent findings on CUBias linked mRNA degradation and/or storage: indeed, AU-rich mRNAs have been found to locate in P-bodies, potentially leading to accumulation of this transcript in the cell ^91^. In addition, the P-body retention of those transcripts reduce their availability to translation and could further explain the reduced protein level for this condition (see discussion below).

### Variation in CUBias and mRNA structure correlate with differences in translation efficiency

Considering all conditions together, our experimental setup allowed us to determine that variation in mRNA levels explains only around 45% of the variation in protein levels, which fits well previous descriptions in the literature for a wide diversity of experimental systems ^72,92,93^. Such relatively weak explanatory power would not be expected if all mRNAs were translated at a constant rate, and has thereby motivated studies to elucidate which factors are involved in the regulation of translation ^94^. Indeed, the literature suggests that in general variations at the mRNA level do not suffice to predict variation at the protein level ^95^, and that this lack of correlation is worse at the single-cell level than at the cell population level ^96^. Here, we provide evidence that co-variation between mRNA levels and protein levels depends on CUBias of our focal gene. Particularly, the AT-rich shble#3 version displayed the highest mRNA levels but contrasting low amounts of the corresponding protein. A possible explanation for this phenomenon could be the selective translation impairment of AT-rich transcripts. As mentioned above, this can result from P-body retention, which physically sequesters AT-rich mRNAs in cell granules making them unavailable for translation ^91^. Other mechanisms may additionally be involved in selective translation impairment. For instance, in human cells, the protein Schlafen11 has been shown to prevent translation of AU-rich transcripts ^97,98^. Given that the AT-rich shble#4 version displays only a moderate translation impairment, we interpret that the dramatic phenotype of shble#3 (high mRNA levels and low protein levels) arises in fact from the combination of suboptimal variables for which a role in optimizing the expression of heterologous genes had already been evidenced ^11^ : (i) similarity to human average CUBias; (ii) the CpG frequency; and (iii) the mRNA folding energy.

i. gene versions with a better match to the average CUBias result in higher protein-to-mRNA ratios. This result is in disagreement with previous reports, as well as with descriptions showing the very limited impact of CUBias on gene expression in mammals, compared to other features ^30,99^. Nevertheless, it is complicated to disentangle the effect of CUBias from other composition characteristics, such as GC and GC3 content. It is even more difficult to interpret them in terms of neutralist or selectionist origin, as both evolutionary hypotheses could account for variation in either parameter (10).
ii. Regarding intragenic CpG frequency, we report a negative impact of very low CpG values on translation efficiency. Such direct effect of low CpG values on translation efficiency had never been reported before, and CpG frequency had been shown to impact heterologous protein amount through its impact on *de novo* transcription instead ^100,101^. More precisely, high CpG depletion was previously associated to low mRNA levels, that weren’t evidenced as a result of changes in nuclear export, alternative splicing or mRNA stability ^100,101^. Indeed, a signature for selection towards decreased values of CpG has been consistently reported ^102,103^, experimentally verified by the detrimental effects of increased CpG levels on protein synthesis ^104,105^, and further corroborated through experimental evolution ^106^.
iii. Regarding the total mRNA folding energy, we also report a negative impact on translation of extreme values. Molecular modelling, along with experimental studies, suggests a prominent role of the initiation steps, rather than elongation steps, on the translation efficacy ^41,107,108^. And indeed, several studies addressing the impact of mRNA folding on translation, established the importance of the 5’ mRNA secondary structure in translation initiation. A shared trend has been identified in bacteria, yeast, protists, and mammals ^31,55,108–111^: a reduced mRNA stability near the site of translation initiation is correlated to a higher protein production. In bacteria and yeast, strong folding around the start codon prevents ribosome recruitment ^31,108^; and a “ramp” of rare codons along the 50 to 100 first coding nucleotides has been reported, with the effect of reducing mRNA folding energy and with the proposed consequences of avoiding ribosome traffic jam ^111,112^. A systematic exploration using 244,000 synthetic DNA sequences on *E. coli* has shown that variation in secondary mRNA structure stability immediately around the start codon accounts for around 36% of the total variance in protein synthesis, while variation in downstream mRNA folding energy accounts only for *ca*. 4-5% ^113^. Nonetheless, an important role of translation elongation cannot be ruled out. Particularly, in human transcripts, de Sousa Abreu and coworkers describe no effect of the initiation rate on translation efficiency ^92^. A recent study in human cell lines, highlights the consequences of the secondary structures along the CDS in the functional half life of mRNA ^110^, which can be related to overall GC and GC3 content as well as to CUBias ^89,90^. In our experimental setup all constructs share by design the nucleotide sequence around the start codon: the 5’UTR corresponds to the plasmid backbone and the first 24 coding nucleotides are identical (AU1 tag). Thus, there are actually no differences in folding energy when considering only the immediate sequence stretch around the start codon, but there are instead differences when considering the full mRNA length. We interpret that our observation of a non-monotonic effect of the full-length mRNA folding energy on the protein-to-mRNA ratio is related to translation elongation impairment rather than to an effect on translation initiation.

### Variation in CUBias modifies intensity and distribution of the fluorescent reporter

We have analysed the fluorescence pattern of the cell populations by means of cytometry. We report phenotypic variability of transfected human cells, observable as multimodal distribution of cellular fluorescence. The multimodal distribution of cellular fluorescence intensity on the transfected cells could be captured in all cases by fitting to a combination of two Gaussian curves. This pattern was similar for all constructs expressing *egfp*, including the empty control and we interpret that it reflects phenotypic plasticity and may be related to transient cellular states, such as cell division status and/or differential kinetics of recovery from transfection-induced cellular stress. Similar differences in gene expression have been actually reported when using cytomegalovirus-based expression vectors ^114^, and have been proposed to be related to cell-cycle dependent cellular states. This bimodal pattern is notwithstanding puzzling, and deserves more attention using a tailored experimental design, that our setup cannot provide. Beyond the shared bimodal distribution of fluorescence levels, we observe significant and concerted shifts of both cellular subpopulations towards higher (*e*.*g*. for the constructs enriched in the most used codons) or lower (*e*.*g*. for constructs using AT-rich codons) values of fluorescence intensity. Thus, differences in overall EGFP-based fluorescence between recoded constructs do not arise from differences in the number of positive cells expressing a given quantity of EGFP, but rather from differences in EGFP synthesis at the individual cell level. Our experimental model using human cells shows that CUBias exerts an important effect on the overall levels but also in the cell-based levels of the heterologous protein produced.

### Heterologous gene expression leads to a trade-off between the benefit conferred through antibiotic resistance and the burden imposed by extra protein synthesis

Strong heterologous expression imposes an enormous basal burden on the cellular economy ^115^. This impact on cellular economy is consistent with the broad literature about the direct (*cis*) and indirect (*trans*) costs of translation ^81^: first, because translation is the per-unit most expensive step during biological information flow ^116^, consuming *ca*. 45% of the whole energy supply in human cells in culture ^117^; second, because virtually all ribosomes are bound to mRNA molecules and potentially engaged in translation ^117^, so that highly-transcribed heterologous mRNA increase overall ribosome demand and cause loss of opportunity for cellular gene translation; and third, because heterologous protein synthesis can lead to additional downstream costs by protein folding, protein degradation and off-target effects of mistranslated proteins ^45,118–120^. Additionally, the mismatch between the CUBias of the heterologous gene and of the expression machinery can display strong trans-effects on the cellular homeostasis, by sequestering ribosomes onto mRNAs that hardly progress over translation but also by creating a competition for the tRNA pools ^31,121^. Scarcity of the less common tRNAs is actually a severe limiting factor for protein synthesis in bacteria ^122^, and this pressure over rare tRNAs can become extreme in conditions of stress, or changes in nutritional status ^10,123,124^.

The *shble* gene that we have used as a base for synonymous recoding encodes for a small protein that confers resistance to bleomycin through antibiotic sequestering ^125^. This protein-antibiotic binding is equimolecular and reversible: an SHBLE protein dimer binds two bleomycin molecules ^83^. The strength of the antibiotic resistance conferred is probably a monotonic, function of the SHBLE amount produced. However, our results suggest that the benefit conferred by SHBLE synthesis in the presence of antibiotic is largely exceeded by the cost and burden of heterologous protein synthesis. We show an important trade-off between the intensity of heterologous SHBLE+EGFP protein synthesis and the actual bleomycin resistance levels, consistent with the strong burden on cell economy imposed by heterologous gene overexpression. This cost can be partly levered when ablating EGFP synthesis in the heterologous constructs, so that a substantial fraction of the burden is removed. Overall we conclude that CUBias of the heterologous gene conferring antibiotic resistance differentially impacts cellular fitness as a function of the differences in heterologous protein synthesis.

## CONCLUSION

The main conundrum for scientists approaching CUBias remains the contrast between, on the one hand, the large and sound body of knowledge showing the strong molecular and cellular impact of gene expression differences arising from CUBias and, on the other hand, the thin evidence for codon usage selection at the organismal level. Under the neutral hypothesis, differences in average genome CUBias can be explained by biochemical biases during DNA synthesis or repair (*e*.*g*. polymerase bias) ^126^; and, in vertebrates, CUBias at the gene level may be shaped by their relative position to isochores (*e*.*g*. alternation between GC-rich and AT-rich stretches along the chromosomes) ^127^. In vertebrates, GC-biased gene conversion mechanisms enhance further such local variations ^126,128,129^. The selective explanation, often referred to as “translational selection”, proposes that different codons may led to differences in gene expression, by changes in alternative splicing patterns, mRNA localisation or stability, translation efficiency, or protein folding ^130^. If such CUBias-induced variation in gene expression were associated with phenotypic variation that results in fitness differences, it would, by definition, be subject to natural selection. Nevertheless, differences in fitness associated with individual synonymous changes seem to be mostly of low magnitude, so that selection may only act effectively in organisms with large population sizes ^131^ such as bacteria ^7^, yeast ^132^, nematodes ^133^, but also in fruit flies ^19,20,134,135^, branchiopods ^136^ and amphibians ^137^. In organisms with small population sizes, such as mammals, and particularly humans, evidences of selection for (or against) certain codons remain nevertheless controversial ^22,138^. In the present manuscript, we have intended to contribute to this debate by exploring the multilevel phenotypic consequences of codon usage differences of heterologous genes in human cells. Our results are consistent with a scenario in which the potential evolutionary forces at play in shaping human CUBias, select for a strict control of mRNA processing (*e*.*g*. splicing, and secondary structure, potentially affecting stability and decay), and that the resulting mRNA properties *in fine* impact translation elongation. Notwithstanding, the disparity between predictions and findings encountered in powerful, codon-usage related experimental evolution approaches highlights the gap in our understanding at connecting phenotype and fitness over different integration levels: molecules, cells, tissues and organisms. Despite, or thanks to, the immense body of knowledge accumulated over the last fifty years, the quest for interpreting and integrating the riddle of CUBias over broad scales of time and biological complexity remains tempting and unsolved.

## MATERIAL AND METHODS

### Design of the *shble* synonymous versions and plasmid constructs

In the present work we have used as focal gene the bleomycin resistance gene present in the genome of the actinobacterium *Streptoalloteichus hindustanus* (ATCC 31158, GenBank X52869.1). We have chosen to focus on this *shble* gene for a number of reasons: 1) because of the mechanism of action of the antibiotic: bleomycin is cytotoxic by intercalating and introducing breaks in the dsDNA ^66^. In this experimental setup we are interested in mRNA translation process, and many antibiotics interfere with protein synthesis at different levels, so that we chose a focal gene with no impact on the mechanisms that we will be evaluating. 2) because of the mechanism that confers resistance: the SHBLE protein interacts on an equimolecular fashion with the bleomycin antibiotic, so that the SHBLE homodimer binds and sequesters two bleomycin molecules ^125^, without performing any catabolic activity on them nor on any other cellular metabolite. The antibiotic resistance level conferred is thus expected to be a direct, monotonic function of the total amount of SHBLE protein produced. 3) because of the small size of the protein synthesised: the SHBLE protein is barely 124 amino acids long, thus minimising the total length of the heterologous mRNA and the impact of translation on the host cell. We did not consider the use of the wild type *shble* sequence in *S. hindustanus* as a meaningful control in our experimental setting using mammalian cells in culture, and we have thus focused on recoding strategies to maximise synonymous differences with respect to our human cell expression system. Six synonymous versions of the *shble* gene were designed applying the “one amino acid--one codon” approach, *i*.*e*., all instances of one amino acid in the *shble* sequence were recoded with the same codon, as follows (Table 1): shble#1 used the most frequent codons in the human genome; shble#2 used the GC-richest among the two most frequent codons; shble#3 used the AT-richest among the two most frequent codons; shble#4 used the least frequent codons; shble#5 used the GC-richest among the two less frequent codons; and shble#6 used the AT-richest among the two less frequent codons. An invariable *AU1* sequence was added as N-terminal tag (amino acid sequence MDTYRI) to all six versions (Sup. Fig. 1).

Nucleotide content between versions are compared in Sup. Table 1. Our recoding strategy succeeds at maximising differences in nucleotide content and to explore a large sequence space in terms of total GC, GC3 and transcript folding energy (Table 1; Sup. Fig. 2). The normalized COUSIN 18 score (COdon Usage Similarity Index), which compares the CUBias of a query against a reference, was calculated using the online tool (http://cousin.ird.fr) ^68^. A score value below 0 informs that the CUBias of the query sequence is opposite to the reference CUBias; a value close to 1 informs that the query CUBias is similar to the reference CUBias, and a value above 1 informs that the query CUBias is similar the reference CUBias, but of larger magnitude ^68^. All *shble* synonymous sequences were chemically synthesised and cloned on the *XhoI* restriction site in the pcDNA3.1+P2A-EGFP plasmid (InvitroGen), in-frame with the *P2A-EGFP* reporter cassette. In this plasmid, the expression of the reporter gene is located under the control of the strong human cytomegalovirus (CMV) promoter and terminated by the bovine growth hormone polyadenylation signal. All constructs encode for a 1,602 bp transcript, encompassing a 1,182 bp *au1-shble-P2A-EGFP* coding sequence (Sup. Fig. 1). The folding energies of all 1,602 bp transcripts were calculated using the RNAfold Webserver (http://rna.tbi.univie.ac.at/cgi-bin/RNAWebSuite/RNAfold.cgi) ^69^, with default parameters (Table 1). During translation, the P2A peptide (sequence NPGP) induces ribosome skipping ^67^, meaning that the ribosome does not perform the Gly-Pro transpeptidation bond and releases instead the AU1-SHBLE moiety and continues translation of the EGFP moiety. The HEK293 human cell line used here is proficient at performing ribosome skipping on the P2A peptide ^139^ The transcript encodes thus for one single coding sequence but translation results in the production of two proteins: SHBLE (theoretical molecular mass 17.2 kDa) and EGFP (27.0 kDa). As controls we used two plasmids: (i) pcDNA3.1+P2A-EGFP (named here “empty”), which encodes for the EGFP protein; (ii) pcDNA3.1+ (named here “superempty”) which does not express any transcript from the CMV promoter (Table 1). In order to explore the burden of EGFP expression we generated two additional constructs by subcloning the AU1-tagged shble#1 and shble#4 coding sequences in the XhoI restriction site of the pcDNA3.1+ backbone, resulting in the constructs shble#1* and shble#4*, lacking the *P2A-EGFP* sequence.

### Transfection and differential cell sampling

All experiments were carried out on HEK293 cells (ACCT CRL-1573). Cell culture conditions, transfection methods and related reagents are detailed in Sup. Methods 2.2. Cells were harvested two days after transfection and submitted to analyses at four levels (Figure 7): (i) nucleic acid analyses (qPCR and RNAseq); (ii) proteomics (label-free quantitative mass spectrometry analysis and western blot immuno-assays); (iii) flow cytometry; and (iv) real-time cell growth analysis (RTCA). Overall, the different experiments were performed on 33 biological replicates, corresponding to a variable number of repetitions depending on the considered analysis (Sup. Method 1). Transfection efficiency was evaluated by means of qPCR targeting two invariable regions of the plasmid and revealed no significant differences between the constructs (Sup. Methods 2.3).

**Figure 7.**
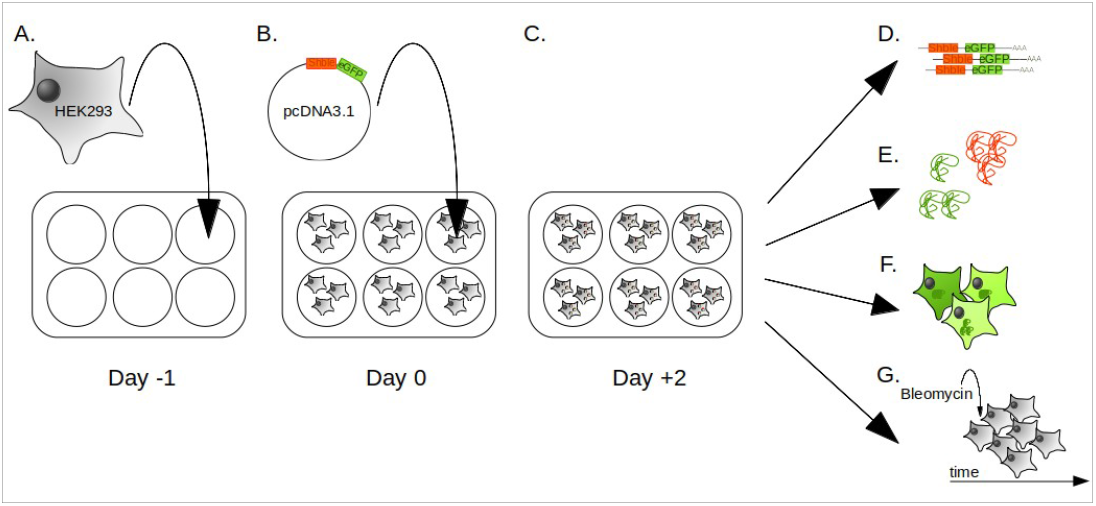
Overview of the sampling protocol and the measured phenotypes. HEK293 cells were seeded on 6-well plates (A) one day before transfection with the customized pcDNA3.1 plasmids (B). Transfected cells were harvested two days later (C). mRNA levels were assessed by RNAseq (D), protein levels were measured by label-free proteomics (E), EGFP fluorescence was assessed at the single cell level by flow cytometry (F) and cell growth was assessed by xCELLigence RTCA (Real Time Cell growth Analysis) in presence of different concentrations of the bleomycin antibiotic (G).

### RNA sequencing and data analysis

Transcriptomic analysis was performed on six biological replicates and eight conditions: shble#1 to shble#6, #empty, and mock (for which the sample is submitted to the exact same procedures, including the transfection agent, but in absence of plasmid). Paired 150bp Illumina reads were trimmed (Trimmomatic v0.38) ^140^ and mapped on eight different genomic references (HISAT2 v2.1.0) ^141^, corresponding to the concatenation of the human reference genome (GCF_000001405.38_GRCh38.p12_genomic.fna, NCBI database, 7^th^ of February 2019) and the corresponding full sequence of the plasmid. For the mock condition, we considered the human genome and all possible versions of the plasmid. Virtually no read of those negative controls mapped to the plasmid sequences. For all other conditions, read distribution patterns along the plasmid sequence were evaluated with IGVtool ^142^. In all cases the *au1-shble-p2a-EGFP* coding sequence displayed highly similar coverage shape for all constructs, except for shble#4 and shble#6 for which respectively one and two alternative splicing events were observed (Sup. Fig. 3 and 4). None of these splice sites were predicted when the theoretical transcripts were evaluated using *Human Splicing Finder* (HSF, accessed via https://www.genomnis.com/access-hsf) ^70^, or with *SPLM - Search for human potential splice sites using weight matrices* (accessed via http://www.softberry.com/) ^71^. When relevant, the three alternative transcript isoforms identified were further used as reference for read pseudomapping and quantification with Kallisto (v0.43.1) ^143^. Details on RNA preparation and bioinformatic pipeline are provided in Sup. Methods 2.4 and Sup. Methods 3.

### Label-free proteomic analysis

Label-free proteomic was performed on nine biological replicates (three of them measured independently, and six pooled by two), and eight different conditions: shble#1 to shble#6, #empty, and mock. For each sample, 20 to 30 µg of proteins were digested in-gel and the resulting peptides were analysed online using a Q Exactive HF mass spectrometer coupled with an Ultimate 3000 RSLC system (Thermo Fisher Scientific). MS/MS analyses were performed using the Maxquant software (v1.5.5.1) ^144^. All MS/MS spectra were searched by the Andromeda search engine ^145^ against a decoy database consisting in a combination of *Homo sapiens* entries from Reference Proteome (UP000005640, release 2019_02, https://www.uniprot.org/), a database with classical contaminants, and the sequences of interest (SHBLE and EGFP). After excluding the usual contaminants, we obtained a final set of 4,302 proteins detected at least once in one of the samples. Intensity based absolute quantification (iBAQ) values were used to compare protein levels between samples ^146^.

### Western blot immunoassays and semi-quantitative analysis

Western blot immunoassays were performed on nine replicates and nine conditions: shble#1 to shble#6, #empty, #superempty, and mock. Three different proteins were targetted: β-TUBULIN, EGFP, and SHBLE (*via* the invariable AU1 epitope tag). Analysis from enzyme chemoluminiscence data was performed with ImageJ ^147^ by «plotting lanes» to obtain relative density plots (Sup. Fig. 7).

### Flow cytometry analysis

Flow cytometry experiments were performed on a NovoCyte flow cytometer system (ACEA biosciences). 50,000 ungated events were acquired with the NovoExpress software, and further filtering of debris and doublets was performed in R with an in-house script (filtering strategy is detailed in Sup. Method 2.7). For subsequent analysis, 30,000 events were randomly picked up from each sample. Seven samples had less than 30,000 viable events and, in order to ensure the same sample size for all conditions, the four corresponding replicates were excluded. After a first visualization of the data, two replicates were ruled out because they displayed a typical pattern of failed transfection for the condition shble#1 (Sup. Method 2.7), resulting in 16 final replicates being fully examined.

### Real time cell growth analysis (RTCA)

RTCA was carried out on an xCELLigence system for the mock and the superempty controls, and further eight constructs: the previously analysed shble#1 to shble#6, plus the shble#1* and shble#4* lacking the *EGFP* reporter gene. Cells were grown under different concentrations of the Bleomycin antibiotic ranging from 0 to 5000 μg/mL (Sup. Method 2.8). Three to six biological replicates were performed, including technical duplicates for each replicate. Cells were grown on microtiter plates with interdigitated gold electrodes that allow to estimate cell density by means of impedance measurement. Measures were acquired every 15 minutes, over 70 hours (280 time points). Impedance measurements are reported as “Cell Index” values, which are compared to the initial baseline values to estimate changes in cellular performance linked to the expression of the different constructs (Sup. Figure 16). For each construct we estimated first cellular fitness by calculating the area below the curve for the delta-Cell index *vs* time for the cells grown in the absence of antibiotics. We estimated then the ability to resist the antibiotic conferred by each construct through calculation of IC50 as the bleomycin concentration that reduces the area below the curve to half of the one estimated in the absence of antibiotics (detailed methods in Sup. Method 2.8).

### Data availability

RNAseq raw reads were deposited on the NCBI-SRA database under the BioProject number PRJNA753061. The mass spectrometry proteomics data have been deposited to the ProteomeXchange Consortium via the PRIDE ^148^ partner repository with the dataset identifier PXD038324.

## ACKNOWLEDGEMENTS

This study was supported by the European Union’s Horizon 2020 research and innovation program under the grant agreement CODOVIREVOL (ERC-2014-CoG-647916) to I.G.B. We acknowledge the IRD itrop HPC (South Green Platform) at IRD Montpellier for providing HPC resources that have contributed to the research results reported within this paper. We also acknowledge the facilities of the Functional Proteomics Platform (FPP) of the Proteomics Pole of Montpellier (PPM, Montpellier France); and the MRI imaging facility, member of the France-BioImaging national infrastructure supported by the French National Research Agency (ANR-10-INBS-04, «Investments for the future»).

