## supplementary file for "Transcriptomic, proteomic and functional consequences of codon usage bias in human cells during heterologous gene expression"

### SUPPLEMENTARY MATERIALS

#### List of supplementary figures

Supplementary figure 1. Customized region of the plasmid vector.

Supplementary figure 2. Principal component analysis of the six shble gene versions, based on four composition variables.

Supplementary figure 3. RNA read distribution along the plasmid sequence, viewed on the Integrative Genomics Viewer (IGV) tool.

Supplementary figure 4. Alternative isoforms and putative splice acceptor sites, viewed on the Integrative Genomics Viewer (IGV) tool.

Supplementary figure 5. Lack of correlation between the transcript representation (total heterologous mRNA) and the plasmid quantity (DNA level).

Supplementary figure 6. Western blot experiments: semi-quantitative analysis of SHBLE and EGFP protein levels, and the ratio between them.

Supplementary figure 7. Western blot and corresponding semi-quantitative analysis with ImageJ.

Supplementary figure 8. Original picture of the western blot membrane shown in Sup. Fig 6.

Supplementary figure 9. Pearson's correlation of SHBLE protein level and *shble* transcript level.

Supplementary figure 10. Pearson's correlation between the total fluorescence signal and the estimated level of EGFP protein.

Supplementary figure 11. Fluorescence distribution per construct and replicate.

Supplementary figure 12. Fluorescence signal distribution: Gaussian Mixture Model (GMM) and its diagnostics for the positive control.

Supplementary figure 13. Positive correlation between the total signal of fluorescence and its central value.

Supplementary figure 14. Principal component analysis based on fluorescence analysis.

Supplementary figure 15. Total fluorescence *vs* cell fitness with or without antibiotics.

Supplementary figure 16. Evolution of the delta Cell Index (CI) over time, depending on the construct and the bleomycin concentration.

#### List of supplementary tables

Supplementary table 1. Number of shared nucleotides (in green) and percentage of identity (in blue) between the six 390 bp au1-shble synonymous sequences.

Supplementary table 2. mRNA abundance: medians of the TPM values for the three alternative transcripts, and ratios of full-to-total mRNA levels.

Supplementary table 3. Protein abundance: median of the normalized iBAQ values for SHBLE and EGFP, and ratios of SHBLE-to-EGFP protein levels.

Supplementary table 4. SHBLE-to-*shble* (protein-to-mRNA) ratio.

---

#### List of supplementary methods

Supplementary method 1. Description of the biological sampling, replicates, and performed experiments.

Supplementary method 2. Details on methods.

Supplementary method 3. Command lines for the RNAseq analysis.

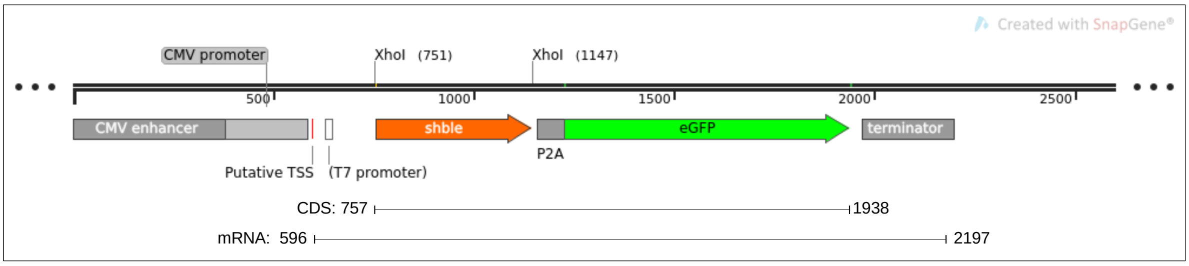


**Supplementary figure 1. Customized region of the plasmid vector.** From 5' to 3', the different sequences come as follows: the CMV (cytomegalo virus) enhancer and promoter were adjacent and located between the position #0 (arbitrarily defined origin) and position #584 base pairs (bp) (in grey). The putative TSS (transcriptional start site) was previously determined at #596 bp (http://www.molecularlab.it/Public/data/GFPina/200691394312_pcdna3.1_man.pdf). The *shble* CDS coupled to the *au1* tag (in orange), were inserted between #757 and #1,146 bp, followed by the *P2A* sequence between #1,159 and #1,224 bp (in grey) and the *egpf* reporter gene between #1,458 and #2,172 bp (in green). The BGH (bovine growth hormone) polyadenylation signal and transcript terminator sequence were located between #1,970 and 2,197 bp (in grey).

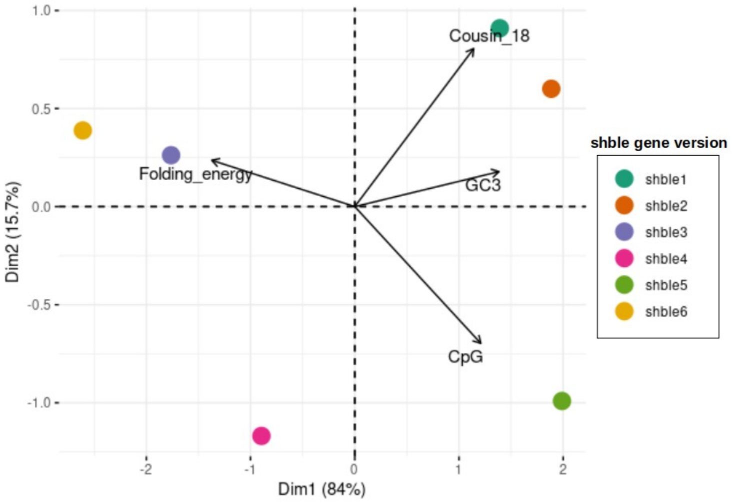


**Supplementary figure 2. Principal component analysis of the six *shble* gene versions, based on four composition variables.** The six versions are: most common #1 (dark green), common and GC-rich #2 (orange), common and AT-rich #3 (purple), rarest #4 (pink), rare and GC-rich #5 (light green), rare and AT-rich #6 (yellow). The four plotted composition variables are: COUSIN (for COdon Usage Similarity INdex – calculated with the online tool http://cousin.ird.fr); the percentage in GC3 content; the percentage in CpG dinucleotides; and the folding energy (i.e. the free energy of the full-length 1,602 bp transcript, calculated on the RNAfold Webserver). Values are given in Table 1.

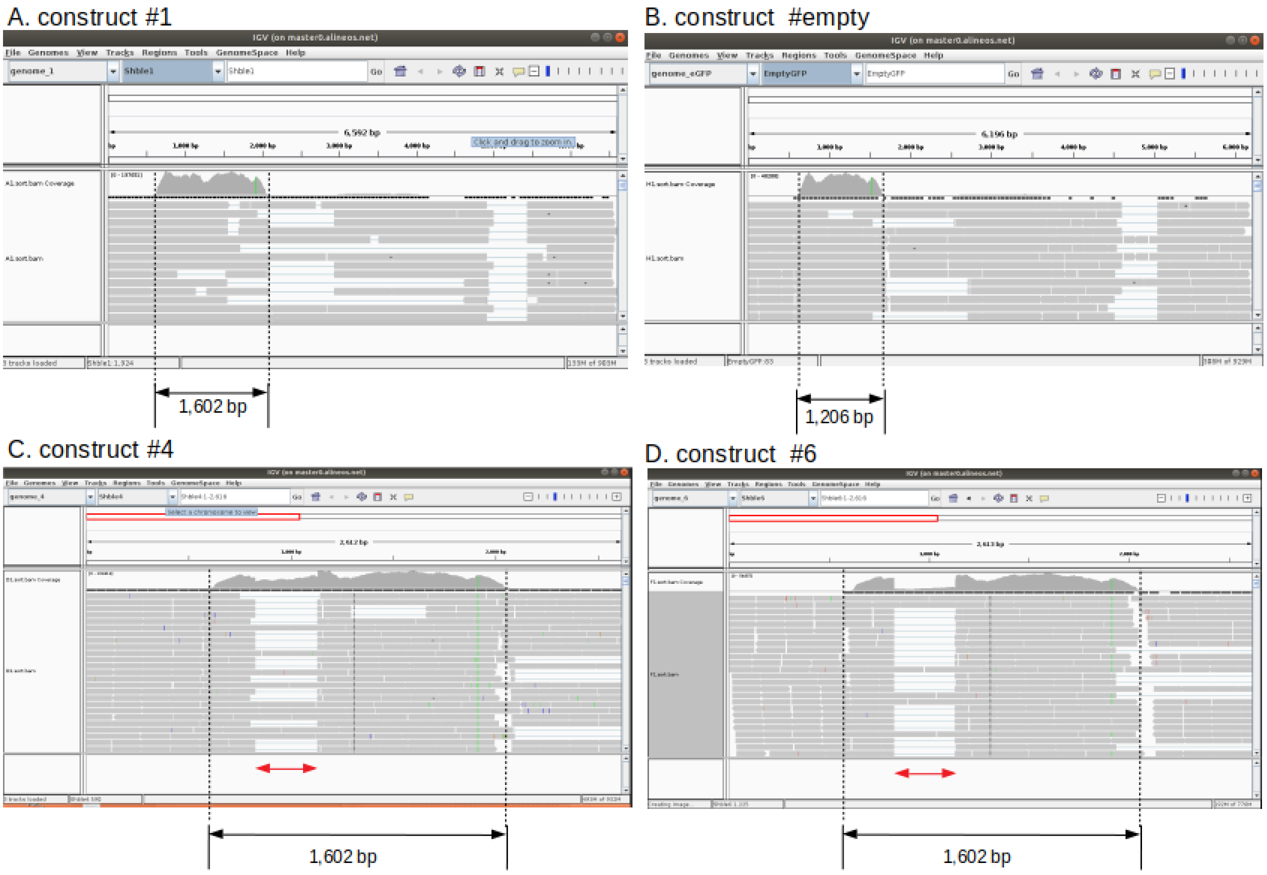


**Supplementary figure 3. RNA read distribution along the plasmid sequence, viewed on the Integrative Genomics Viewer (IGV) tool.** Typical observed patterns for the plasmid sequence are: a distribution of the RNA reads through a region of 1,602 base pairs for all constructs containing both the *shble* and the *EGFP* genes (Panel A, black arrow); or reads along a 1,206 bp region for the #empty construct (Panel B, black arrow). An unexpected pattern was observed for two of the constructs (#4 and #6), for which one region was less covered (red arrow) within the typical 1,602 bp sequence (Panels C and D).

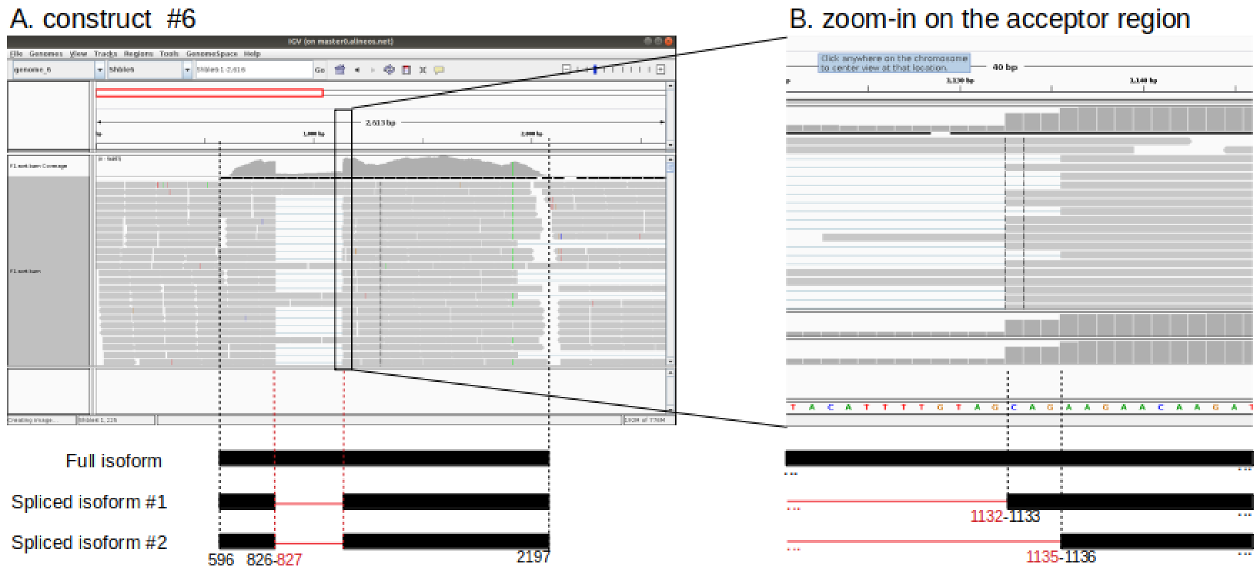


**Supplementary figure 4. Alternative isoforms and putative splice acceptor sites, viewed on the Integrative Genomics Viewer (IGV) tool.** Panel A: The typical covered region began at base #596 - base #0 being the first base of the CMV - and ended at base #2,197. It corresponds to the full isoform of 1,602 bp. For shble#6, the region that was covered by less reads began at base #827 (CAG/GTG) (Panel A, in red) and ended alternatively at base #1,132 (TAG/CAG) or #1,135 (CAG/AAG) (Panel B, in red), defining two alternative spliced forms. For shble#4, the nucleotide sequence at the donor site is: CGG/GTG and at the acceptor is: TAG/CGG (data not shown).

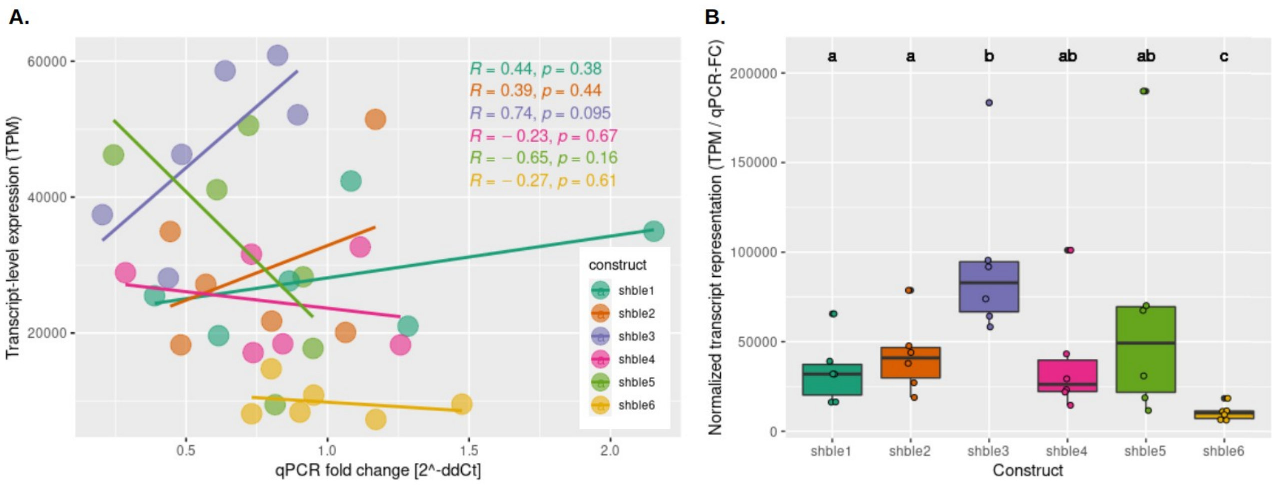


**Supplementary figure 5. Lack of correlation between the transcript representation (total heterologous mRNA) and the plasmid quantity (DNA level).** On both panels, six biological replicates of six different conditions are shown: #1 (dark green), #2 (orange), #3 (purple), #4 (pink), #5 (light green) and #6 (yellow). Panel A: linear regression analysis between the total number of heterologous transcript (TPM) and the plasmid quantity (qPCR fold change). *R* is the Pearson's correlation coefficient and *p* the associated p-value. Panel B: heterologous TPM normalized by the qPCR fold change for each of the six experimental conditions. Statistics are pairwise Wilcoxon rank sum test, with Benjamini-Hochberg adjusted p-values: conditions associated to a same letter do not display different median values of TPM.

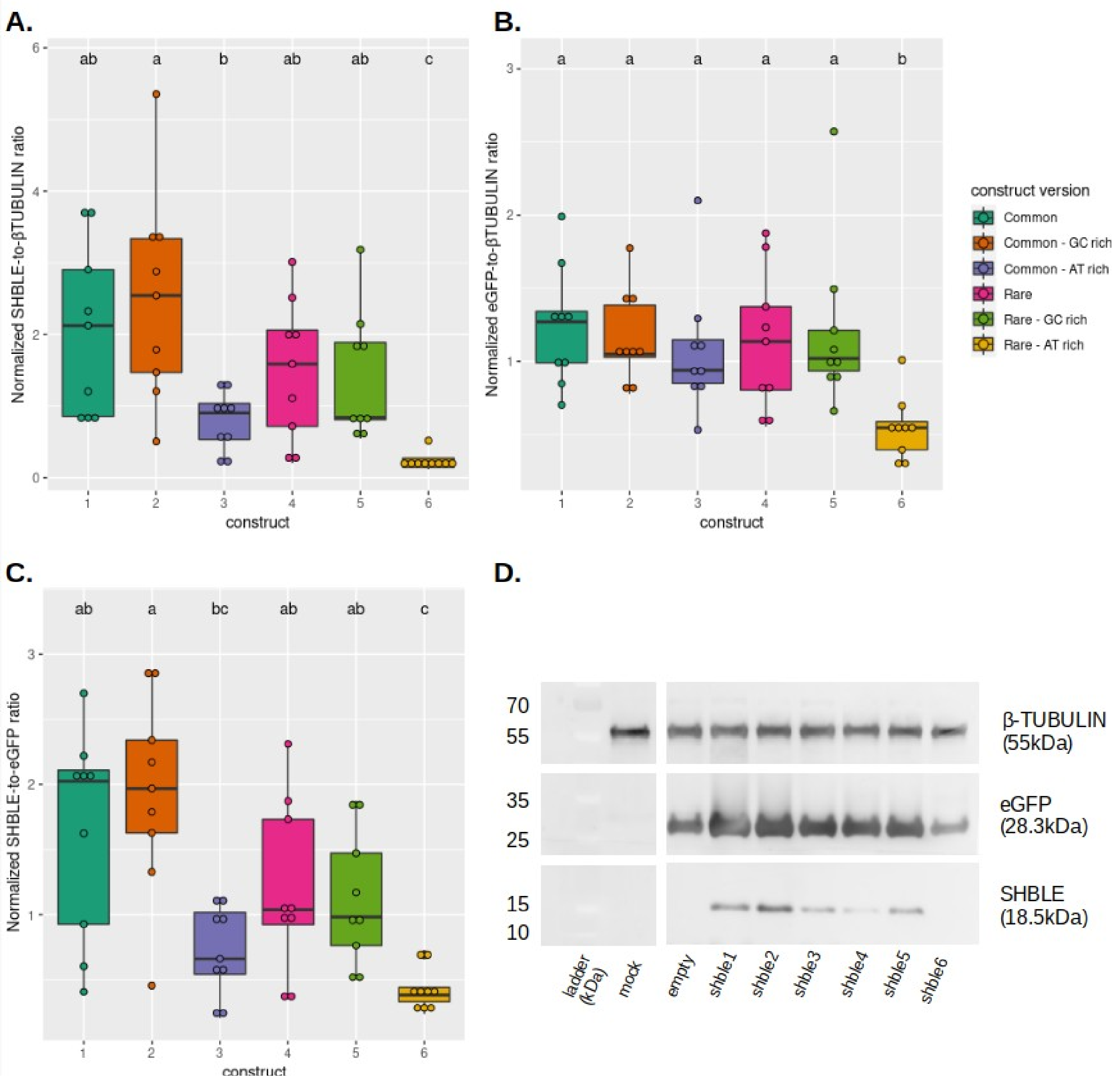


**Supplementary figure 6. Western blot experiments: semi-quantitative analysis of SHBLE and EGFP protein levels, and the ratio between them.** Panel A: estimated SHBLE protein abundance. Panel B: estimated EGFP protein abundance. Panel C: SHBLE-to-EGFP ratio. Values shown on Panel A to C are based on the area below the curve of relative density with automatic baseline (ImajeJ measurement, Sup. Fig 7). Six different conditions, and 9 biological replicate for each, are shown: shble#1 (dark green), shble#2 (orange), shble#3 (purple), shble#4 (pink), shble#5 (light green) and shble#6 (yellow). Statistical test is a pairwise wilcoxon rank sum test, with Benjamini-Hochberg adjusted p-values: conditions associated to a same letter do not display different median values of the corresponding variable. Panel D: Immunoblotting of β-TUBULIN (upper band), EGFP (middle band) and SHBLE (lower band) for eight conditions including the mock (no plasmid) and empty (*EGFP* but no *shble* CDS). The original gel is provided as Sup. Fig 8.

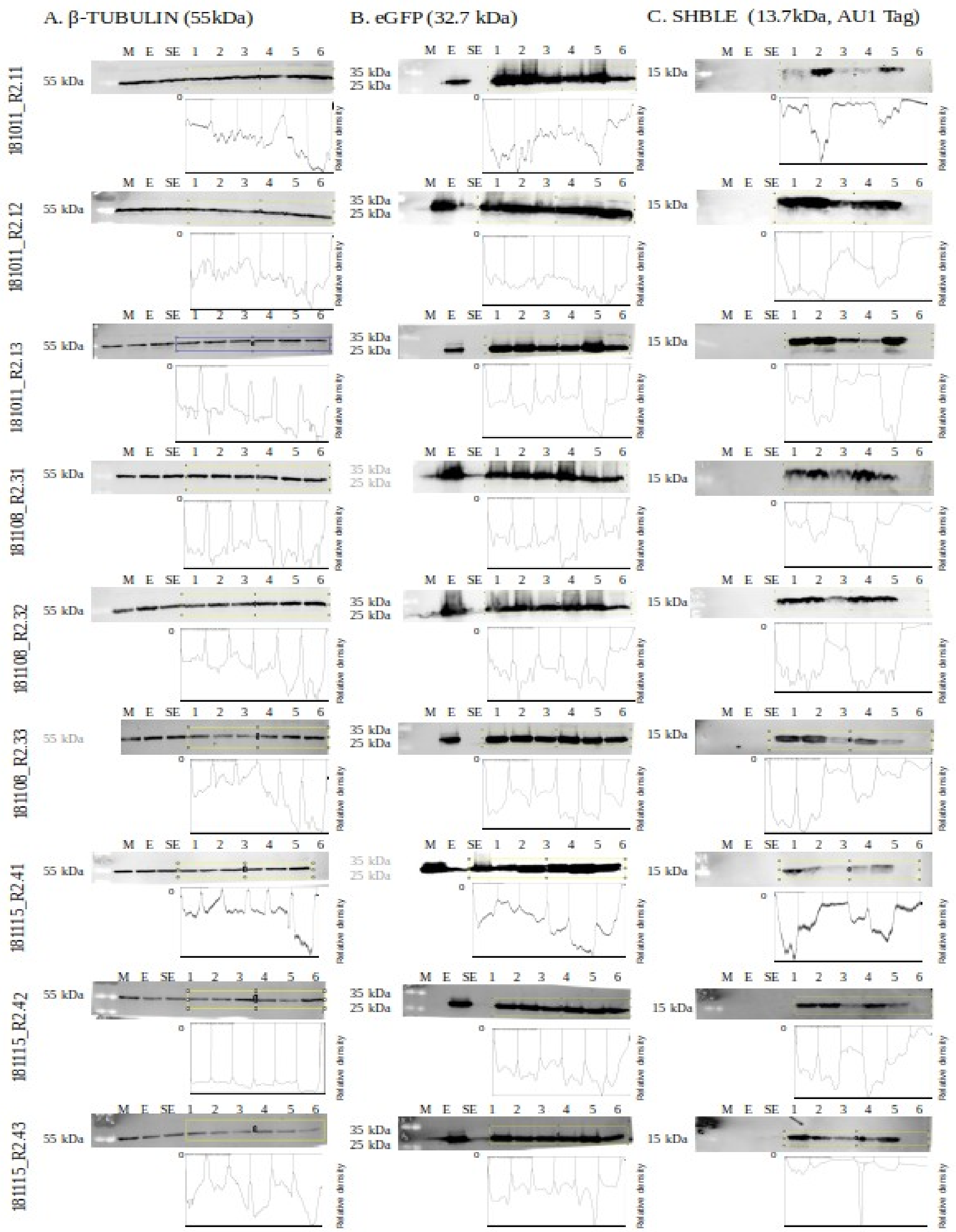


**Supplementary figure 7. Western blot and corresponding semi-quantitative analysis with ImageJ.** One line corresponds to one replicate and one to column corresponds to one targeted protein (β- TUBULIN in A, EGFP in B and SHBLE in C). Below each blot, the curve of relative density obtained with the ImageJ « plotting lanes » function is shown, the automatic baseline is indicated by a black 0 on the left of each curves. Each peak corresponds to a band. Corresponding plots of SHBLE and EGFP estimated protein abundance and the ratio between them are shown in Sup. Fig 6.

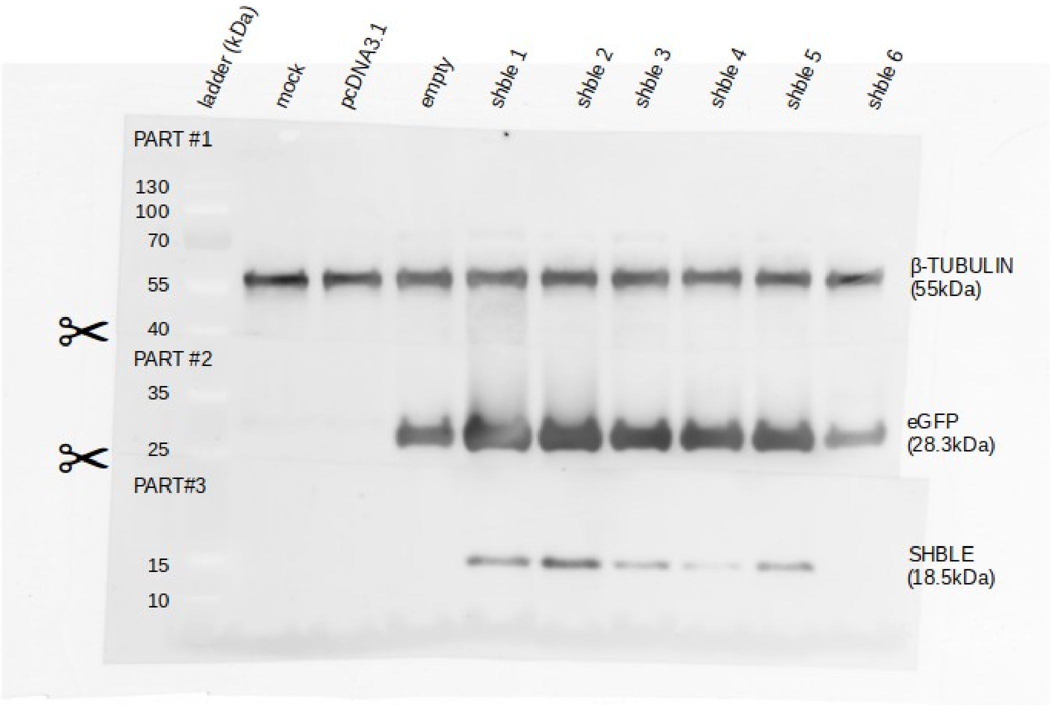


**Supplementary figure 8. Original picture of the western blot membrane shown in Sup. Fig 6.** After electroblotting to nitrocellulose paper, the membrane was cut in three pieces (see scissors symbol), and incubated separately: Part #1 - above 40kDa – antibody anti-βtubulin, concentrated at 1:2000 (Thermo Fisher Scientific, 11305883); Part #2 - between 40 and 25 kDa - antibody anti-EGFP, concentrated at 1:8000 (Origene, TA150032); Part #3 - below 25 kDa – antibody anti-P2A epitope, concentrated at 1:800 (Novus, NBP2-59627), P2A being linked to the upstream SHBLE. Secondary antibody was anti-rabbit, conjugated to horseradish peroxidase (HRP) (656120, Thermo Fisher Scientific, 1:3000). For the image acquisition, the PhotoMultiplier Tube (PMT) was set to 500V.

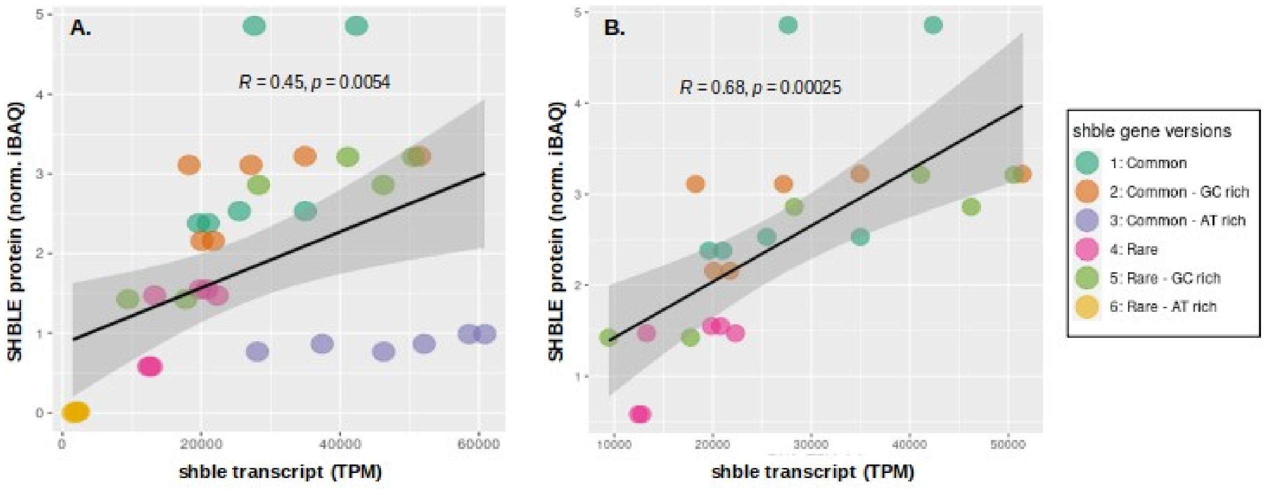


**Supplementary figure 9. Pearson's correlation of SHBLE protein level and *shble* transcript level.** Six different conditions are shown: #1 (dark green), #2 (orange), #3 (purple), #4 (pink), #5 (light green) and #6 (yellow). Panel A: The correlation takes into account all six constructs. Panel B: The constructs #3 and #6, which displayed a discordant pattern from the others (Figure 3A), was excluded from the calculation. Six biological replicates are represented (all have independent RNAseq measures, but were pooled by 2 for the label free analysis).

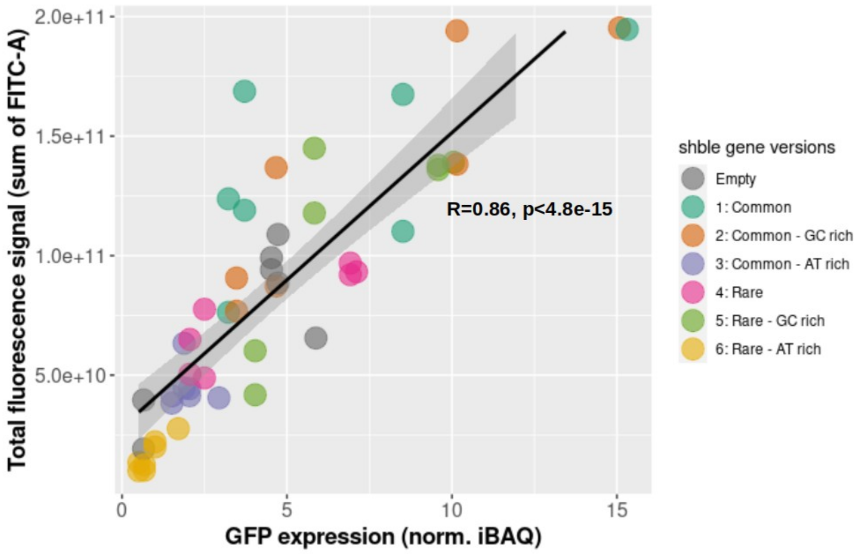


**Supplementary figure 10. Pearson's correlation between the total fluorescence signal and the estimated level of EGFP protein.** Seven different conditions are shown: the positive control « empty » (grey), #1 (dark green), #2 (orange), #3 (purple), #4 (pink), #5 (light green) and #6 (yellow). Seven biological replicates are represented (all have independent fluorescence measures, but 6 were pooled by 2 for the label-free analysis). The total fluorescence signal corresponds to the sum of all fluorescence values (expressed as FITC-A unit). The EGFP protein level is expressed as normalized iBAQ.

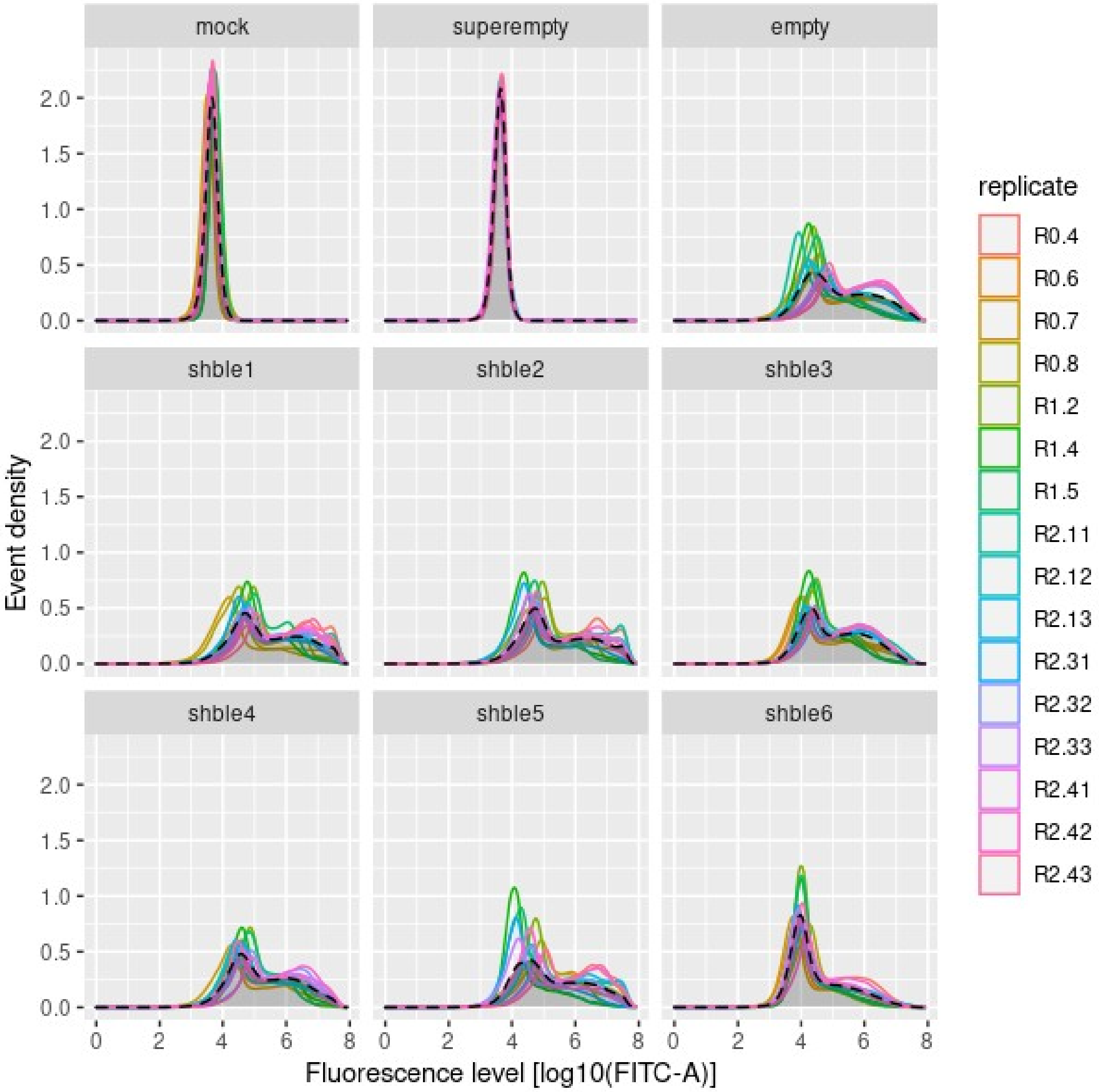


**Supplementary figure 11. Fluorescence distribution per construct and replicate.** Six tested conditions (shble#1 to #6), two negative controls (« mock » without plasmid, and « superempty » with the pcDNA3.1 missing the *au1-shble-P2A-EGFP* part), and one positive control (« empty », missing only the *au1-shble* part) are individually represented per panel. Each color represents one of the 16 replicates (remaining after all the filtering steps), corresponding to 30,000 individual events (i.e. cells) for each condition. The black dashed curve filled in grey corresponds to the distribution of the fluorescence signal considering all the events of one given condition: they are compared in the Figure 4 and Table 2.

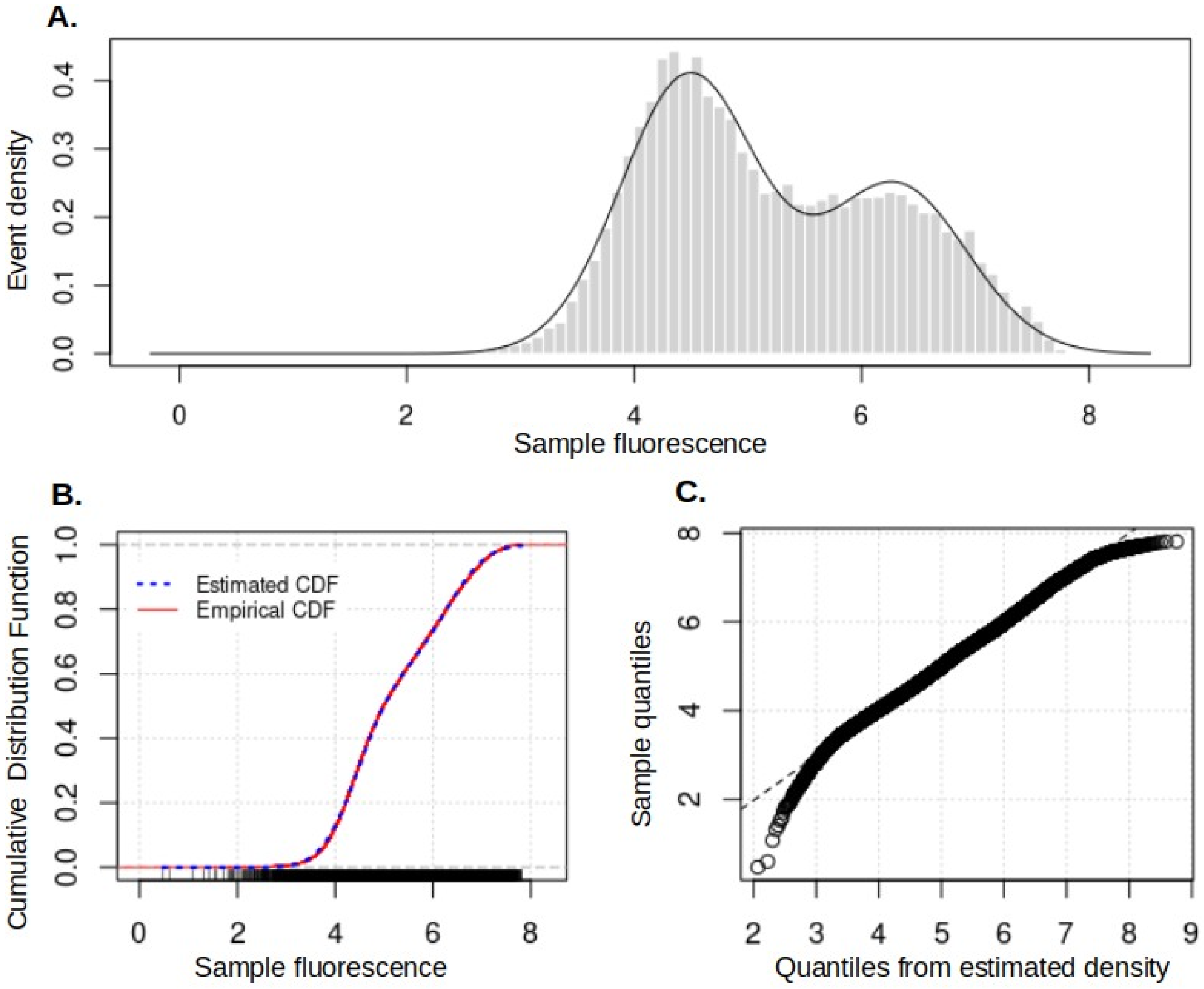


**Supplementary figure 12. Fluorescence signal distribution: Gaussian Mixture Model (GMM) and its diagnostics for the positive control.** The represented sample is the the «empty» construct (i.e. missing *au1-shble* but containing the *P2A-EGFP* region). Panel A represents the fluorescence distribution of a subset of 30,000 events (i.e. randomly picked up among the 16 replicates) (grey bars) and the modeled distribution corresponding to the GMM (black line). Panel B shows the overlap between the estimated "CDF" (Cumulative Distribution Function - dotted blue line) and the empirical CDF (continuous red line). Panel C is a Q-Q plot of the sample quantiles, versus the quantiles obtained from the inverse of the estimated CDF: the points form a line that is roughly straight, meaning that both sets of quantiles come from similar distributions. Thus, the estimated GMM with two Gaussian appear as a good model for the empirical fluorescence signal distribution.

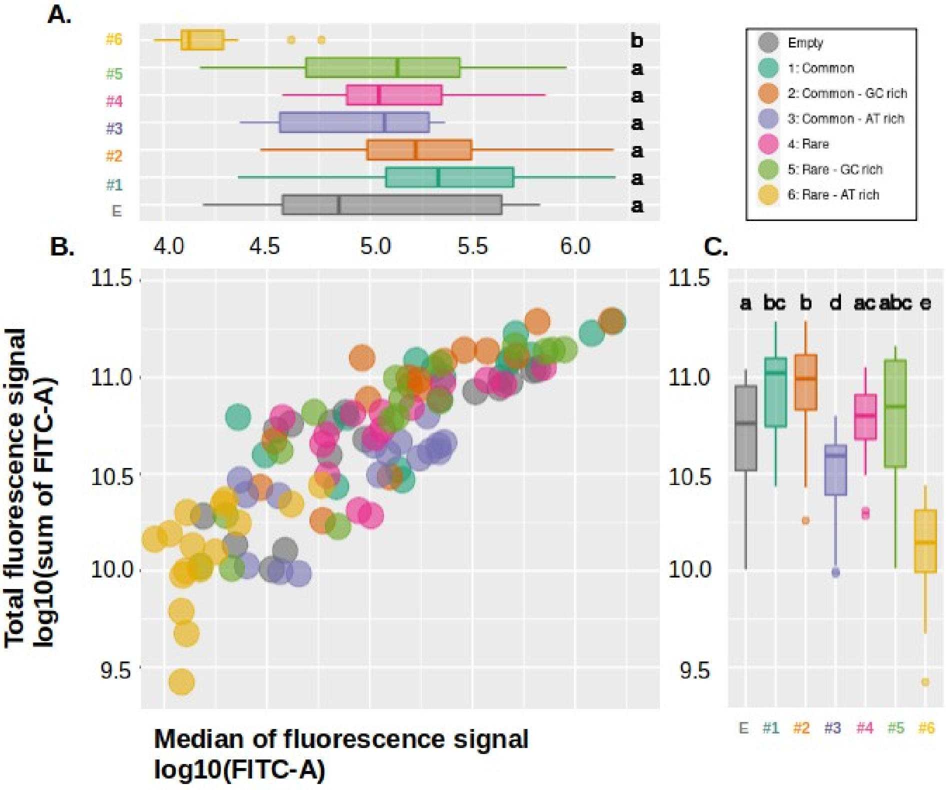


**Supplementary figure 13. Positive correlation between the total signal of fluorescence and its central value.** Sixteen biological replicates and seven different conditions are shown: empty (grey), #1 (dark green), #2 (orange), #3 (purple), #4 (pink), #5 (light green) and #6 (yellow). Marginal boxplots (A) and (C) respectively show the median of the fluorescence signal (as log10(FITC-A) and the total fluorescent signal (as log10 of the sum FITC-A). Statistical test is a pairwise wilcoxon rank sum test, with Benjamini-Hochberg adjusted p-values. Panel B depicts the correlation between the total signal and the median, corresponding Pearson coefficient is R=0.85 (p-value<2.2e-16).

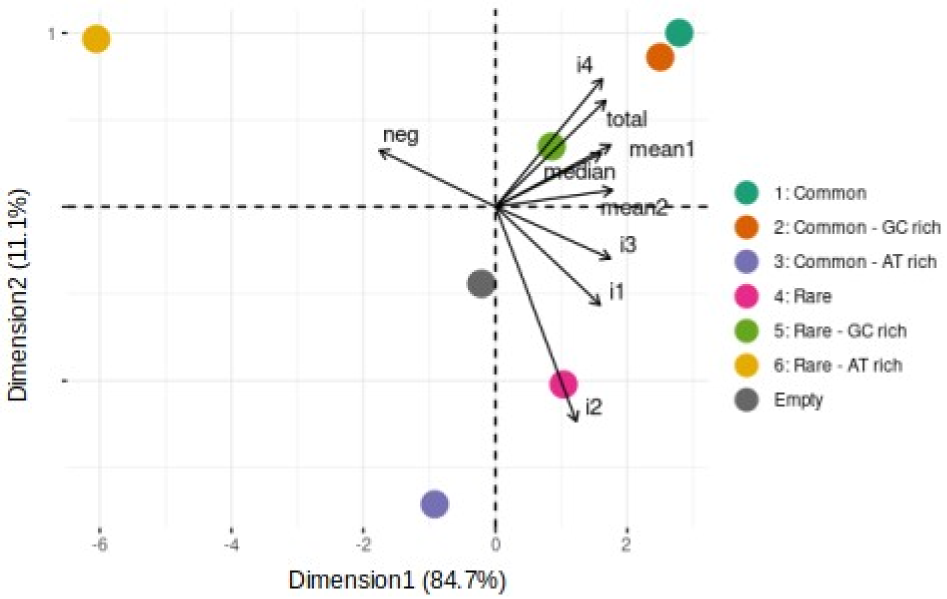


**Supplementary figure 14. Principal component analysis based on fluorescence analysis.** Abbreviations are: « neg » for the number of negative cells, « total » for the sum of all FITC-A, « median » for the median of FITC-A, « mean1 » for the mean of the first underlying Gaussian curve (in log10(FITC-A)), « mean2 » for the mean of the second underlying Gaussian curve, « i1 » to « i4 » for the number of events within each interval.

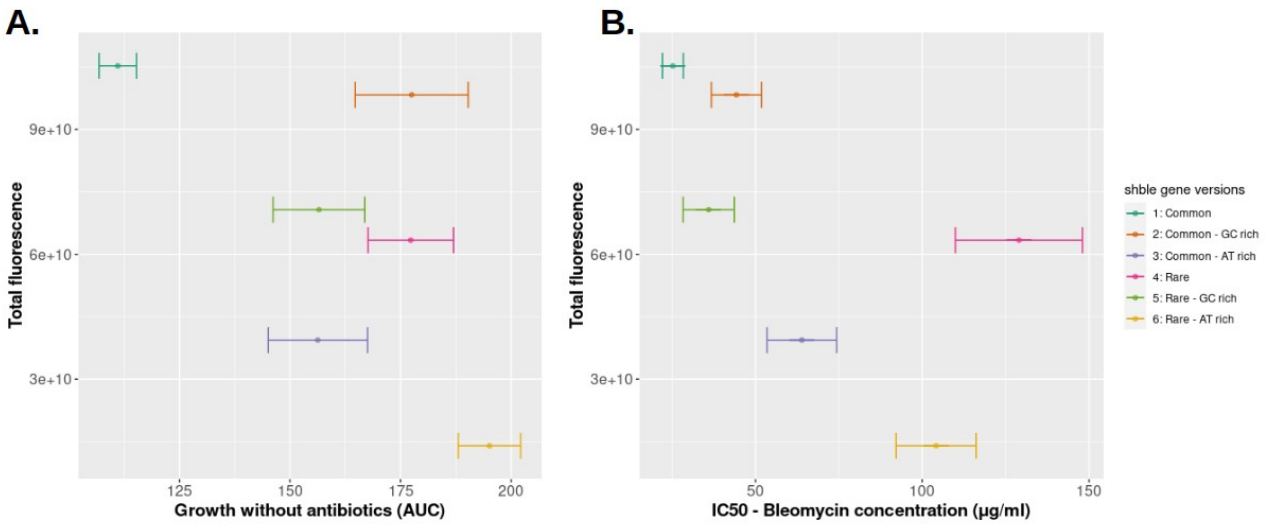


**Supplementary figure 15. Total fluorescence *vs* cell fitness with or without antibiotics.** Panel A : Cell fitness in absence of antibiotics is represented by the maximum cell growth in absence of antibiotics, proxied as the area under the curve of the delta Cell Index ("AUC") ; Panel B : Cell fitness in presence of antibiotics is represented by IC50 (the bleomycin concentration that reduces to 50% the corresponding growth). For x axes of both panels, the plotted central values were estimated fitting variation of Cell Index data to Hill's equation (pooled data, 3 to 6 biological replicates), and bars correspond to the estimated standard error. For y axes, the plotted central values correspond to the median of the total fluorescence signal of 16 biological replicates, and the respective standard error are negligible (plotted but not visible).

Colors correspond to six different experimental conditions : #1 (dark green), #2 (orange), #3 (purple), #4 (pink), #5 (light green), #6 (yellow).

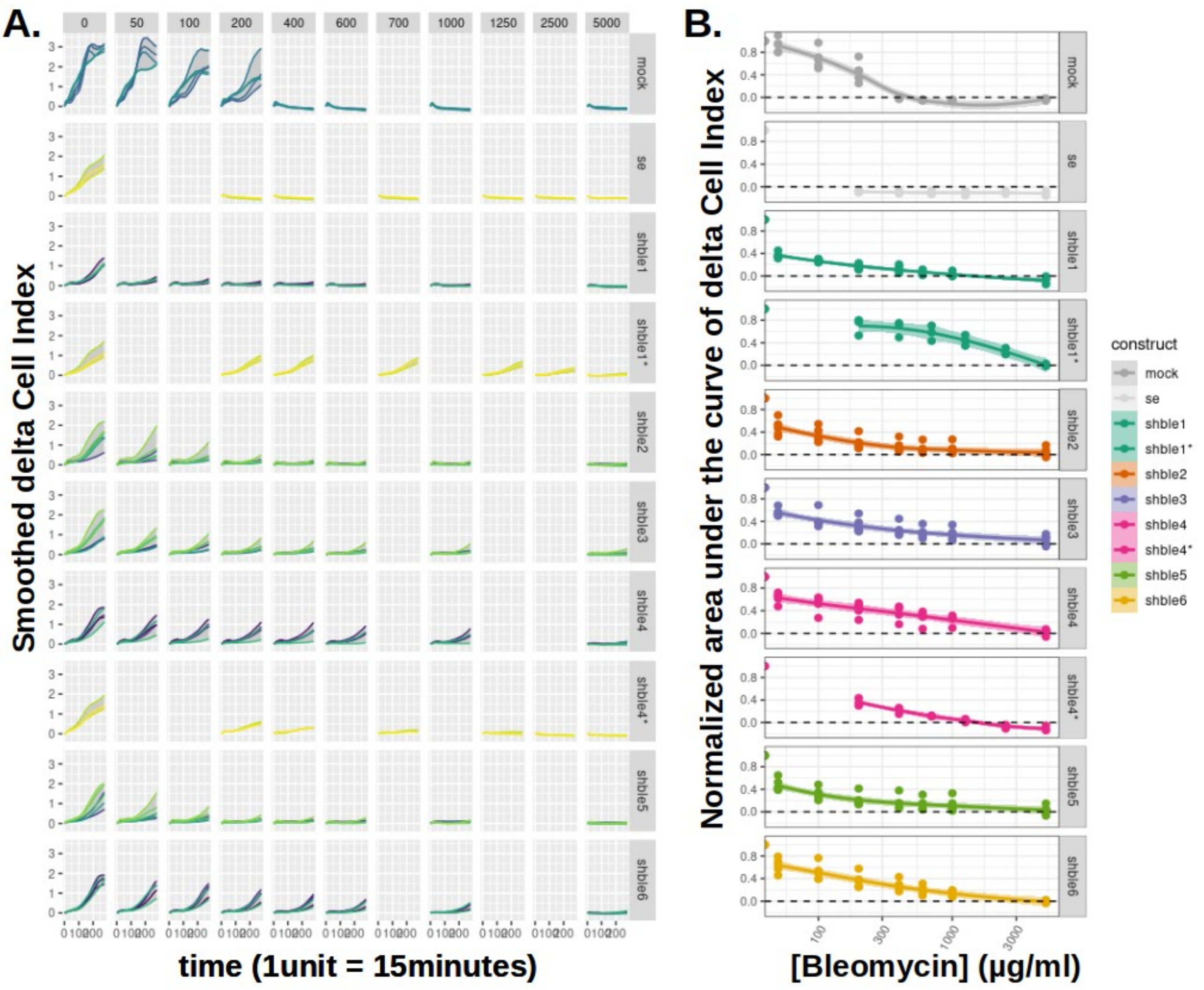


**Supplementary figure 16. Evolution of the delta Cell Index (CI) over time, depending on the construct and the bleomycin concentration.** Panel A: Each row corresponds to one condition, up-to-bottom: mock, superempty, and shble #1 to #6. The 4th and 8th rows respectively correspond to shble#1* and shble#4* lacking the *EGFP* reporter gene. The columns (left-to-right) correspond to increasing concentrations of antibiotics (from 0 to 5000 µg/ml). Measure acquisition was done every 15 minutes during 70 hours (280 time points). The delta CI values correspond to the CI values from which we substracted the reference CI at t=0, within each considered technical replicate. The curves correspond to smoothed delta CI values (loess curve fitting) of technical duplicates (one color per biological replicate). Panel B: Ordinates represent the area under the curve (AUC) of deltaCI, normalized by the AUC in absence of antibiotics. Abscissa represent the different concentration of antibiotics (from 0 to 5000 µg/ml – log scale). Each row corresponds to one condition: mock (darkgrey), superempty (lightgrey), #1 (dark green), #2 (orange), #3 (purple), #4 (pink), #5 (light green), #6 (yellow) and, shown by stars on the 4th and 8th rows, the versions shble#1* (dark green) and shble#4* (pink) lacking the *EGFP* reporter gene.

**Supplementary table 1. Number of shared nucleotides (in green) and percentage of identity (in blue) between the six 390 bp *AU1-shble* synonymous sequences.** The most similar versions are shble#1 and shble#2 (95.90 % of identity), while the versions shble#1 and shble#4 are the most dissimilar (65.04 % of identity). Values were obtained performing a blastn alignment optimizing for “more dissimilar sequences (discontiguous megablast)”.

| | shble#1 | shble#2 | shble#3 | shble#4 | shble#5 | shble#6 |
|---|---|---|---|---|---|---|
| shble#1 | | 95.90 % | 73.78 % | 65.04 % | 73.59 % | 70.18 % |
| shble#2 | 374 | | 69.67 % | 67.10 % | 75.64 % | 66.07 % |
| shble#3 | 287 | 271 | | 75.64 % | 65.55 % | 89.49 % |
| shble#4 | 253 | 261 | 295 | | 80.98 % | 82.56 % |
| shble#5 | 287 | 295 | 255 | 315 | | 65.55 % |
| shble#6 | 273 | 257 | 349 | 322 | 255 | |

**Supplementary table 2. mRNA abundance: medians of the TPM values for the three alternative transcripts, and ratios of full-to-total mRNA levels.** The three first columns are TPM values corresponding to (i) the detected level of full mRNA, (ii) the first and (iii) the second spliced forms. The fourth column gives the sum of TPM values of the three transcript forms. The last column corresponds to the ratio of TPM of the full mRNA over the sum of all three transcripts (i.e. full-to-total).

| | **full** | **isoform 1** | **isoform 2** | **total** | **full-to-total** |
|---|---|---|---|---|---|
| **shble#1** | 26564.70 | 0.36 | 5.02 | 26570.08 | 99.98 % |
| **shble#2** | 24458.90 | 0.53 | 4.10 | 24463.53 | 99.98 % |
| **shble#3** | 49184.25 | 1.10 | 14.00 | 49199.35 | 99.96 % |
| **shble#4** | 16566.15 | 7229.52 | 10.86 | 23806.53 | 69.44 % |
| **shble#5** | 34678.50 | 0.00 | 2.37 | 34680.87 | 99.99 % |
| **shble#6** | 1979.10 | 5008.20 | 1962.46 | 8949.75 | 20.93 % |

**Supplementary table 3. Protein abundance: median of the normalized iBAQ values for SHBLE and EGFP.** Normalized iBAQ corresponds to the iBAQ value over the median of the 100 best iBAQ values within a given sample.

| | **SHBLE** | **EGFP** |
|---|---|---|
| **shble#1** | 6.407964 | 11.91341 |
| **shble#2** | 4.556977 | 10.34301 |
| **shble#3** | 0.9265822 | 2.048282 |
| **shble#4** | 2.060895 | 7.0136 |
| **shble#5** | 3.101033 | 9.812794 |
| **shble#6** | 0.0316293 | 1.353385 |

**Supplementary table 4. SHBLE-to-*shble* (protein-to-mRNA) ratio.** The ratio corresponds to the protein level expressed as normalised iBAQ, divided by the full-length transcript levels expressed as TPM value. Letters refer to the results of a pairwise Wilcoxon rank sum test: conditions associated with a same letter do not display significantly different median values ($p<0.05$ after Benjamini-Hochberg correction).

| | **SHBLE-to-*shble* ratio** | **W. test** |
|---|---|---|
| **shble#1** | 113.99 e-06 | a |
| **shble#2** | 103.29 e-06 | ab |
| **shble#3** | 16.73 e-06 | c |
| **shble#4** | 70.26 e-06 | b |
| **shble#5** | 79.27 e-06 | ab |
| **shble#6** | 6.51 e-06 | d |

### SUPPLEMENTARY METHODS

#### Supplementary method 1. Description of the biological sampling, replicates, and performed experiments.

| | *Nomenclature* | | | *Phenotype* | | *Global approches* | | *Targeted approaches* | |
|---|---|---|---|---|---|---|---|---|---|
| | **Sampling date (yymmdd)** | **Replicate number** | **Name in analyses** | **Xcelligence** | **Flow cytometry** | **Label-free proteomics** | **RNA-seq** | **qPCR** | **Westernblot** |
| Exp #3 | Dec. 2019 | R3.1 | R3.1 | • (se, 1&4 wo gfp) | no | no | no | no | • |
| | Dec. 2019 | R3.2 | R3.2 | • (se, 1&4 wo gfp) | no | no | no | no | • |
| | Dec. 2019 | R3.3 | R3.3 | • (se, 1&4 wo gfp) | no | no | no | no | • |
| Exp #2 | 181011 | R2.11 | 181011_R2.11 | no | • | • (mix R2.11 & R2.13 → R2.1) | • | • | • |
| | 181011 | R2.12 | 181011_R2.12 | no | • | no | no | • | • |
| | 181011 | R2.13 | 181011_R2.13 | no | • | • (mix R2.11 & R2.13 → R2.1) | • | • | • |
| | 181108 | R2.31 | 181108_R2.31 | no | • | no | no | • | • |
| | 181108 | R2.32 | 181108_R2.32 | no | • | • (mix R2.32 & 2.33 → R2.3) | • | • | • |
| | 181108 | R2.33 | 181108_R2.33 | no | • | • (mix R2.32 & 2.33 → R2.3) | • | • | • |
| | 181115 | R2.41 | 181115_R2.41 | no | • | • (mix R2.41 & 2.43 → R2.4) | • | • | • |
| | 181115 | R2.42 | 181115_R2.42 | no | • | no | no | • | • |
| | 181115 | R2.43 | 181115_R2.43 | no | • | • (mix R2.41 & 2.43 → R2.4) | • | • | • |
| Exp #1 | 180615 | R1.1 | 180615_R1.1 | no | • (not analyzed) | | no | no | no |
| | 180622 | R1.2 | 180622_R1.2 | no | • | no | no | no | no |
| | 180629 | R1.3 | 180629_R1.3 | no | • (not analyzed) | no | no | no | no |
| | 180706 | R1.4 | 180706_R1.4 | no | • | no | no | no | no |
| | 180706 | R1.5 | 180706_R1.5 | no | • | no | no | no | no |
| Exp #0 | 171005 | R0.01 | 171005_R0.01 | • (1,4,6 with gfp) | no | no | no | no | no |
| | 171012 | R0.02 | 171012_R0.02 | • (1,4,6 with gfp) | no | no | no | no | no |
| | 171019 | R0.03 | 171019_R0.03 | • (1,4,6 with gfp) | no | no | no | no | no |
| | 171102 | R0.1 | 171102_R0.1 | • (2,3,5 with gfp) | no | no | no | no | no |
| | 171109 | R0.2 | 171109_R0.2 | no | no | • (3 tech.rep.) | no | no | no |
| | 171116 | R0.3 | 171116_R0.3 | • (2,3,5 with gfp) | • (not analyzed) | • | no | no | no |
| | 171123 | R0.4 | 171123_R0.4 | • (1,4,6 with gfp) | • | • | no | no | no |
| | 171130 | R0.5 | 171130_R0.5 | • (1,4,6 with gfp) | • (not analyzed) | no | no | no | no |
| | 171207 | R0.6 | 171207_R0.6 | • (2,3,5 with gfp) | • | no | no | no | no |
| | 171214 | R0.7 | 171214_R0.7 | • (2,3,5 with gfp) | • | no | no | no | no |
| | 180111 | R0.8 | 180111_R0.8 | • (2,3,5 with gfp) | • | no | no | no | no |
| | 180118 | R0.9 | 180118_R0.9 | no | • (not analyzed) | no | no | no | no |
| | 180125 | R0.10 | 180125_R0.10 | • (2,3,5 with gfp) | • (not analyzed) | no | no | no | no |
| | 180420 | R0.11 | 180420_R0.11 | • (mock x 2) | no | no | no | no | no |
| | 180522 | R0.12 | 180522_R0.12 | • (mock) | no | no | no | no | no |
| | 180615 | R0.13 | 180615_R0.13 | • (mock x 2) | no | no | no | no | no |
| | **Sampling date (yymmdd)** | **Replicate number** | **Name in analyses** | **Xcelligence** | **Flow cytometry** | **Label-free proteomics** | **RNA-seq** | **qPCR** | **Westernblot** |
| | *Nomenclature* | | | *Phenotype* | | *Global approches* | | *Targeted approaches* | |

**Supplementary method 2. Details on methods.**

**1. *shble* synonymous versions and plasmid design.** The original *shble* sequence was obtained from the GenBank database (X52869.1, *https://www.ncbi.nlm.nih.gov/nuccore/X52869.1?report=genbank*). pcDNA3.1(+)-based plasmids were synthesized at GenScript (Genscript Corporation, Piscataway, NJ, USA).

**2. Cell culture, transfection and sampling.** HEK293 cells (Human Embryonic Kidney cells, CRL-1573, ATCC) were cultured at 37°C with 5% CO2, in complete medium: MEM (Minimum Essential Medium Earle, M1MEM10K, Eurobio), with 10% FBS (Foetal Bovine Serum, CVFSVF0001, EuroBio) and 1% antibiotics (Penicillin-Streptomycin, 15140122, Fisher scientific). Transfection was done in 6-well plates, with $1.17x10^5$ cells per mL, seeded at day -1. At day 0, the complete medium was changed for MEM 2% FBS. A given plasmid mixed with turbofect transfection reagent (12331863, Fisher scientific) was added in each well. Cells were harvested at day +2 (Trypsin-EDTA, CEZTDA000U, Eurobio), and prepared for four different experiments: (i) nucleic acid analysis: cells were kept in Monarch DNA/RNA protection reagent (T2011, NEB) at -80°C; (ii) proteomics: cells were kept in RIPA buffer (RadioImmunoPrecipitation Assay buffer) and cOmplete™ Protease Inhibitor Cocktail (04693116001, Sigma Aldrich), at -20°C; (iii) cytometry: cells were fixed (phosphate-buffered saline PBS, CS1PBS0001, Eurobio), with 2% paraformaldehyde (PFA, 10630813, Fisher) and conserved in PBS with 0,1% glycine (G5417, Sigma Aldrich) at 4°C in dark; (iv) Real Time Cell Analysis (RTCA) with the xCELLigence device: cells were transferred in dedicated 48-well plates, with 30,000 cells and 100µl of complete medium per well.

**3. Evaluation of the transfection efficiency: DNA extraction and qPCR experiments.** In order to control for the plasmid quantity in transfected cells, we assessed the normalized representation of each construct by a series of qPCR assays. DNA extraction was done with the DNeasy Blood & Tissue Kit (69504, QIAGEN) following manufacturer's recommendations for cultured cells. qPCR assays were done on a LightCycler 96 system (Roche) with the QuantiNova SYBR Green PCR Kit (208054, QIAGEN) and the following cycling conditions: one cycle of activation (95°C, 2 minutes), 40 cycles of quantification (95°C - 5 min. in alternance with 60°C - 10 min.). The plasmid was targeted in two invariable regions, and three housekeeping genes were assessed. List of primers, and result representation for 9 biological replicates are provided below.

**qPCR primer table:**

| Sequence | Target |
|---|---|
| CTGGAGACGTGGAGGAGAAC | p2a sequence between shble and egfp genes |
| GCTTGCCGGTGGTGCAGATG | egfp gene |
| GAGAACCCACTGCTTACTGG | 5'UTR of the shble gene |
| GCCACTGTGCTGGATATCTG | 5'UTR of the shble gene |
| GACAGTCAGCCGCATCTTCT | Housekeeping gene : GAPDH (Glyceraldehyde-3-phosphate dehydrogenase) |
| TTAAAAGCAGCCCTGGTGAC | Housekeeping gene : GAPDH (Glyceraldehyde-3-phosphate dehydrogenase) |
| ACCGTGTTCTTCGACATTGC | Housekeeping gene : PPIA (Peptidylprolyl Isomerase A) |
| GCCTCCACAATATTCATGCC | Housekeeping gene : PPIA (Peptidylprolyl Isomerase A) |
| TCCTCCACTGGTACACAGGC | Housekeeping gene : tubulin |
| CTCCTCTTCGGCCTCCTCAC | Housekeeping gene : tubulin |

A.

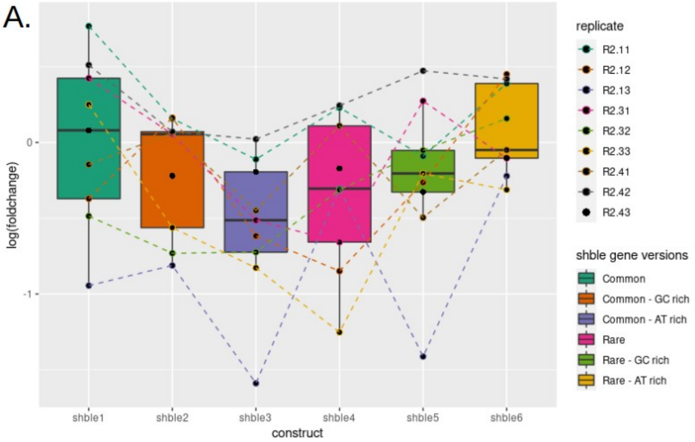


B.

| | shble1 | shble2 | shble3 | shble4 | shble5 |
|---|---|---|---|---|---|
| **shble2** | 0.50 | - | - | - | - |
| **shble3** | 0.30 | 0.41 | - | - | - |
| **shble4** | 0.48 | 0.92 | 0.58 | - | - |
| **shble5** | 0.59 | 0.93 | 0.41 | 0.83 | - |
| **shble6** | 0.93 | 0.50 | 0.06 | 0.39 | 0.48 |

***Normalized plasmid quantity for each experimental condition.*** *Each of the qPCR reaction was performed in technical duplicates and the mean of the two resulting Ct values was used for further analysis. The ΔCt corresponds, for each sample, to the mean of the 3 housekeeping genes substracted from the mean of the 2 plasmid targets: ΔCt=Ct(targets)-Ct(housekeeping genes). For each biological replicate, we then considered the #empty condition as reference, and calculated the ΔΔCt by substracting the ΔCt(empty) from the ΔCt(construct). In A, the plotted quantity correspond to the foldchange=2^-(ΔΔCt) value for each condition. Nine biological replicates are represented. In B, pairwise comparisons between constructs are shown: statistical test is a wilcoxon rank sum test with Benjamini-Hochberg correction. It was performed on ΔΔCt values and reveal no significant differences.*

**4. RNA extraction, sequencing and data analysis**. RNA extraction was done with the Monarch Total RNA miniprep kit (T2010S, NEB), following manufacturer's recommendations. PolyA selection, strand-specific RNA library preparation and 2x150bp sequencing were performed at the Genewiz NGS laboratory (New Jersey, USA) on an Illumina HiSeq4000 instrument. Quality check, trimming and adapter removal were performed respectively with FastQC (v0.11.5, https://www.bioinformatics.babraham.ac.uk/projects/fastqc/) and Trimmomatic (v0.38) (Bolger et al., 2014). The trimmed reads were then mapped on the genomic reference using HISAT2 (v2.1.0) (Kim et al., 2015). We observed an overall alignment rate above 97.5 % for all libraries. The resulting BAM files, sorted by coordinates (samtools v1.3.1) (Li et al., 2009), were used to visualize the plasmid coverage on the Integrative Genomic Viewer (IGV) tool (Robinson et al., 2011). Read pseudomapping and quantification were performed with Kallisto (v0.43.1) (Bray et al., 2016). The human reference transcriptome was obtained at https://www.ncbi.nlm.nih.gov/genome/?term=txid9606[orgn] (GCF_000001405.39_GRCh38.p13_rna.fna, 11th of September 2019). The bioinformatic pipeline is illustrated below and the corresponding command lines are provided as Sup. Method 3.

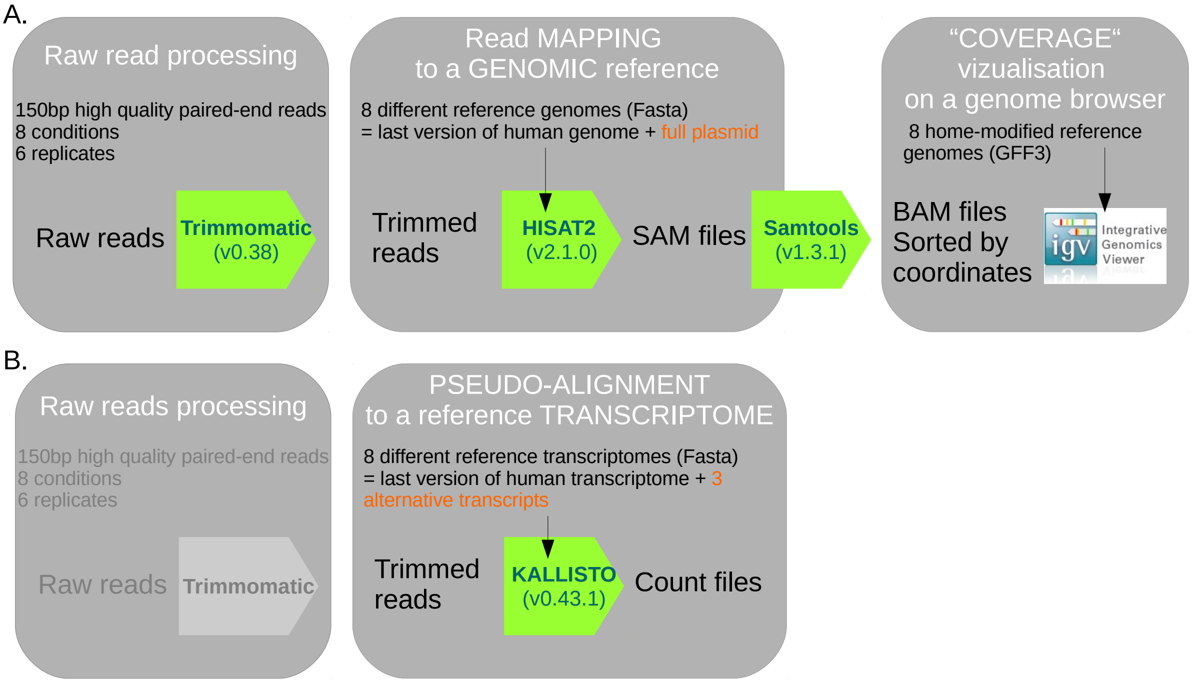


***Bioinformatic pipeline for the RNAseq analysis.*** *RNAseq data were processed in two times: first, the trimmed reads were mapped to the reference genomes in order to observe their distribution along the plasmid (A); second, the trimmed reads were pseudo-mapped to the reference transcriptomes in order to quantify each distinct transcript (B).*

**5. Protein extraction and label-free proteomic analysis.** Mass spectrometry experiments were carried out using facilities of the Functional Proteomics Platform (FPP) of the Proteomics Pole of Montpellier (PPM, Montpellier France). Solubilized proteins were mechanically sheared, and resuspended in Laemmli buffer. 20 to 30 µg of proteins were in-gel digested using Trypsin (Trypsin Gold, Promega), as previously described (Shevchenko et al., 2007). Resulting peptides were resuspended in loading buffer and proteins were analyzed online using Q Exactive HF mass spectrometer coupled with Ultimate 3000 RSLC (Thermo Fisher Scientific) fitted with a stainless-steel emitter (Thermo Fisher Scientific). A gradient consisting of 2-40% B in 123 min (A: 0.1% formic acid in water; B: 0.1% formic acid 80% ACN) at 300 nL/min was used to elute peptides from the capillary reverse-phase column (0.075 mm x 500 mm, Pepmap® C18, Thermo Fisher Scientific). MS/MS analyses were performed in a data-dependent mode with standard settings. Analysis was performed using the Maxquant software (v1.5.5.1) (Cox & Mann, 2008). All MS/MS spectra were searched by the Andromeda search engine (Cox et al., 2011) against a decoy database consisting in a combination of *Homo sapiens* entries from Reference Proteome (UP000005640, release 2019_02, https://www.uniprot.org/), a database with classical contaminants, and the sequences of interest (SHBLE and EGFP). Default search parameters were used, Oxidation (Met) and Acetylation (N-term) as variable modifications and Carbamidomethyl (Cys) as fixed modification were applied. FDR was set to 1% for peptides and proteins. A representative ID in each protein group was automatically selected using an in-house bioinformatics tool (Leading_v3.2). From the 4,734 identified proteins, we excluded the usual contaminants and the proteins with a peptide counts (razor+unique) lower than two, and obtained a final set of 4,302 proteins.

**6. Western blot immunoassays and semi-quantitative analysis**. We used SDS-PAGE 12% gels. After electroblotting, nitrocellulose membranes were cut in order to incubate separately the relevant part with the antibody of interest: β-TUBULIN (Thermofisher Scientific, 11305883, concentration 1:1000), or EGFP (Origene, TA150032, 1:2000 to 1:6000), AU1 epitope tag (Novus, NB600-59627, 1:500 to 1:900), or P2A epitope tag (Novus, NBP2-59627, 1:800. Blots were blocked with 5% non-fat dry milk (Biorad, 170-6404) in phosphate buffered saline with Tween (PBST). Antibody hybridations were performed in PBST-1.7% bovine serum albumin (BSA) at 4°C overnight, or one hour at room temperature. Membranes were then washed with PBST and incubated with anti-rabbit secondary antibody conjugated to horseradish peroxidase (HRP) (656120, Thermo Fisher Scientific, 1:3000). Chemiluminescent detection was performed using ECL prime Western-blotting system (GERPN2232, Sigma Aldrich) and images were aquired with TYPHOON FLA 9500 biomolecular imager (450V to 500V PhotoMultiplier Tube (PMT), and 25 or 50μm pixel size).

**7. Flow cytometry analysis.** Data acquisition was done with the NovoExpress software, with the following parameters: 50,000 ungated events, fast flowrate, treshold FSCH larger than 100.00, excitation at 487nm (FITC-GFP), and acquisition of 600 events per second. Event gating was performed in R with an in-house script, available on the github repository dedicated to this study. Illustrated filtering steps are shown for one sample below. Gating was achieved as follow: (i) First step was to remove the outliers in terms of size and complexity, by excluding the 0.5 % maximum and minimum values for Forward SCatter Height (FSC.H, cell size), Side SCatter Height (SSC.H, complexity), Forward SCatter Area (FSC.A) and Side SCatter Area (SSC.A). (ii) Second step was to remove the debris that display lower size and complexity (first peak of the FSC.H and SSC.H distributions, see figure below). (iii) Finally, doublets were removed by excluding extreme SSC.H/SSC.A ratios. For subsequent analysis, 30,000 events were randomly picked up from each sample, and seven samples with less than 30,000 events were excluded (together with the full replicate they correspond to). After a first visualisation of the data, two replicates were excluded because they displayed a typical pattern of failed transfection for the condition #1 (see figure below). Hence, the results shown in this manuscript are based on 16 full repetitions corresponding to 480,000 events per condition. To compare the distribution of the fluorescence signal between conditions, the Anderson-Darling test was performed with the « k-Samples » R package with default parameters, using 100,000 events randomly sampled among the 16 replicates of each conditions. Besides, the density distribution was described thanks to an Univariate Gaussian Mixture Model (GMM) (Mclust, Sup. Fig. 12). We fixed the number of mixture components to « 2 », and the type of model to be fitted to « V » (i.e. unabling unequal variance).

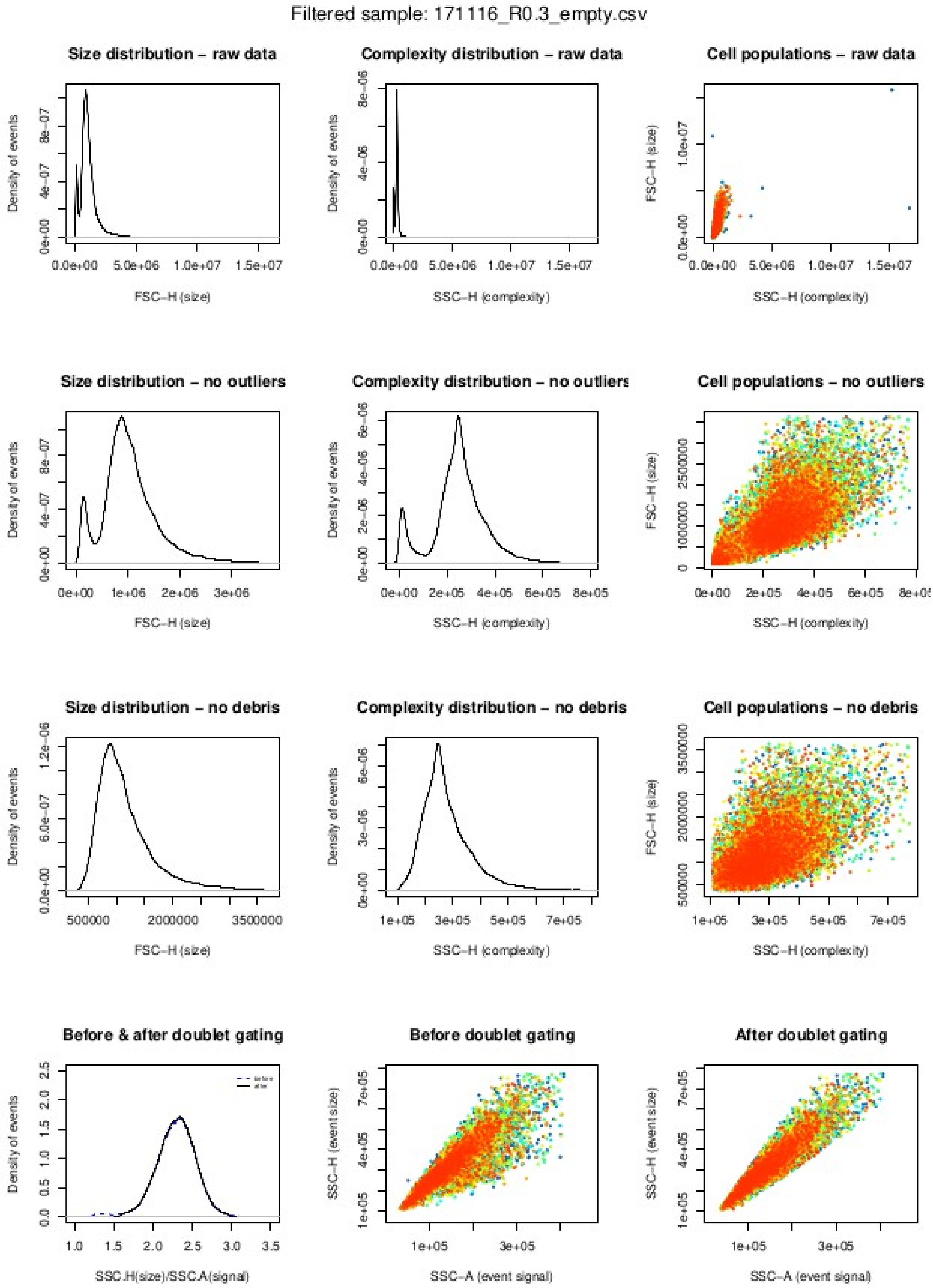


***Strategy of debris and doubet filtering: example of one sample.***

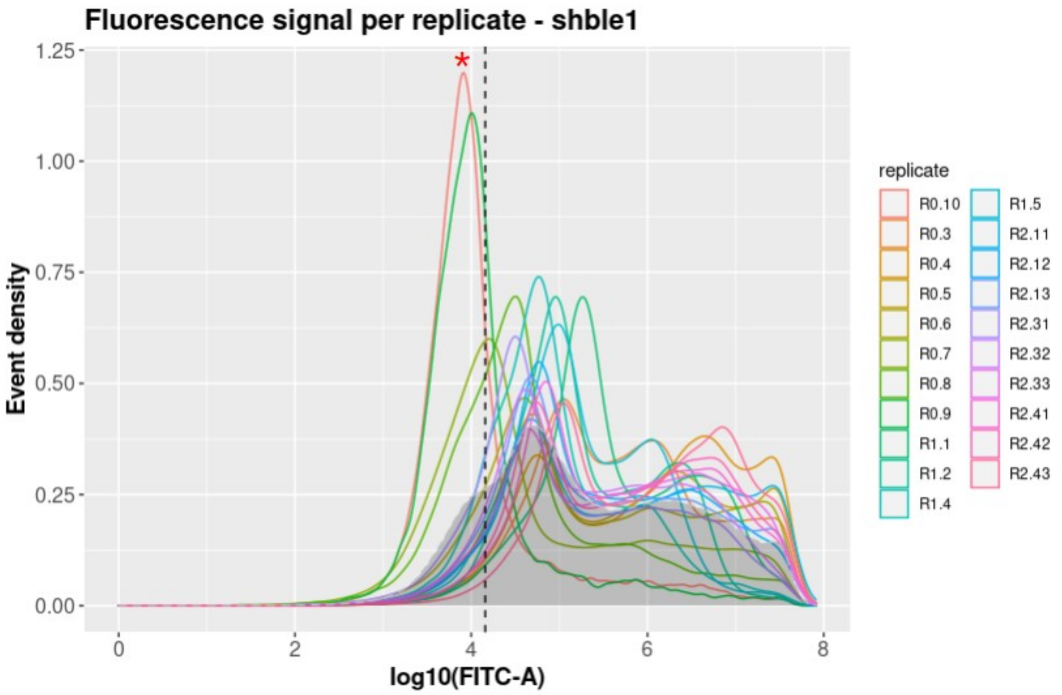


***Distribution of the fluorescence signal for 21 replicates of the shble #1 condition.*** *Each color represents one replicate (see Sup. Method 1 for details on nomenclature). The vertical black dashed line represents the limit of cell positivity (log10 (14486) = 4.160948). Two replicates, R0.10 (in red) and R0.9 (in green), shown by a red star, display a narrow distribution, shifted on the left, characteristic of samples with low level of transfection and not consistent with the 19 other replicates. They were removed from further analysis.*

**8. Real time cell analysis.** This serie of experiments was performed on a xCELLigence RTCA system (Agilent technologies), on two controls (the mock and the « superempty » - *i.e.* pcDNA3.1 plasmid without *shble* neither *EGFP* sequences) and 8 constructs (shble #1 to #6, plus #1* and #4* lacking the *EGFP* reporter gene), in technical duplicate for all conditions, but with variable numbers of biological replicates: 6 rep. (for #2, #3 and #5), 5 rep. (for #1, #4, #6 and mock), or 3 rep. (for the superempty, #1* and #4*). Different concentrations of the Zeocin antibiotic (a member of the bleomycin family, R250-01, Invitrogen) were considered. For the mock and the constructs #1 to #6, eight concentrations were tested: 0, 50, 100, 200, 400, 600, 1000 and 5000 µg/ml; for the superempty, #1* and #4*, seven concentrations were tested: 0, 200, 400, 700, 1250, 2500 and 5000 µg/ml (Sup. Fig. 17). Briefly, 48 hours after transfection, cells were transferred from the 6-well plates to the xCELLigence RTCA 48-well plates, with 30,000 cells and 100µl of complete medium (EMEM, 10% FBS and 1% penicillin-streptomycin) per well. Cells were then incubated at 37°C and 5% $CO_2$. Measures were acquired every 15 minutes, throughout 70 hours (280 time points): the xCELLigence system gives Cell Index (CI) values, a dimensionless parameter, that reflects the biological condition of the monitored cells. This takes into account the cell number, the cell viability, the morphology and the adhesion degree. By definition, if there is no cell or if they are not adhered, CI=0; while for cells having the same physiological condition, the more cells there are, the higher is the CI. In order to account for little variations in the initial cell numbers, we further considered the delta CI, using CI at t=0 as reference. As technical replicates were very consistent, they were merged and the resulting curves were smoothed by a Loess curve fitting (*i.e.* local polynomial regression, see Sup. Fig 17). For each condition and biological replicates, the area under the curve (AUC) of those smoothed delta CI were calculated. In order to take into account the possible differences in a given population of cells, we further considered the fold-change AUC which corresponds to the AUC, normalized by the AUC in absence of antibiotic (Sup. Fig 17). Using those fold-change AUC, IC50 have been calculated with the JMP software (v16), modeling the experimental set using fits to Hill's equation (considering all replicates together). Finally, the IC50 were fixed to fit the Hill's equation back to the raw AUC values, and one unique estimation of the AUC in absence of antibiotic was done for each construct.

**Supplementary method 3. Command lines for the RNAseq analysis.**

##### 1. FastQC (v0.11.5)

```
fastqc path_to_input/all_reads.fastq.gz -o  path_to_output
```

##### 2. Trimmomatic (v0.38)

```
java -jar Trimmomatic-0.38/trimmomatic-0.38.jar PE -phred33
path_to_input/forward_reads.fastq.gz path_to_input/reverse_reads.fastq.gz
path_to_output/forward_reads_paired.fastq.gz
path_to_output/forward_reads_unpaired.fastq.gz
path_to_output/reverse_reads_paired.fastq.gz
path_to_output/reverse_reads_unpaired.fastq.gz
ILLUMINACLIP:Trimmomatic-0.38/adapters/TruSeq3-PE.fa:2:30:10 HEADCROP:13 LEADING:3
TRAILING:3 SLIDINGWINDOW:4:15 MINLEN:100
```

##### 3. FastQC (v0.11.5)

```
# (replace * by 'forward' or 'reverse')
fastqc path_to_input/*_reads_paired.fastq.gz -o path_to_output
```

##### 4. Hisat2 (v2.1.0)

```
# build genome index (replace * by the version of the reference)
hisat2-2.1.0/hisat2-build path_to_input/genome_* RefGenomeIndex
# read mapping
hisat2-2.1.0/hisat2 -strand -p 2 --dta -x path_to_input/RefGenomeIndex -1
path_to_input/forward_reads_paired.fastq.gz -2
path_to_input/reverse_reads_paired.fastq.gz -S library_name.sam
```

##### 5. Samtools (v1.3.1)

```
samtools view -b library_name.sam | samtools sort -o library_name.sort.bam
```

##### 6. Kallisto (v0.43.1)

```
# build transcriptome index (replace * by the version of the reference)
kallisto index -i transcripts.idx path_to_input/transcriptome_*
# read mapping
kallisto quant -t 16 -i path_to_input/transcripts.idx -o path_to_input/library_name -b
100 --fr-stranded path_to_input/forward_reads_paired.fastq.gz
path_to_input/reverse_reads_paired.fastq.gz
```
